# Revealing interspecies transmission barriers of avian influenza A viruses

**DOI:** 10.1101/2020.11.17.386755

**Authors:** Mahmoud M. Naguib, Per Eriksson, Elinor Jax, Jonas Nilsson, Carina Sihlbom, Cecilia Lindskog, Caroline Bröjer, Britt-Marie Olsson, Michelle Wille, Jonas Waldenström, Göran Larson, Robert H. S. Kraus, Åke Lundkvist, Björn Olsen, Josef D. Järhult, Patrik Ellström

## Abstract

Influenza A virus (IAV) pandemics result from interspecies transmission events within the avian reservoir and further to mammals including humans. Investigating molecular virus–host interactions dictating this process and the adaptations to the new hosts that follow is vital to understand zoonotic IAV spread. Receptor incompatibility has been suggested to limit zoonotic IAV transmission from the wild bird reservoir. Other barriers to interspecies transmission, particularly within the avian system, largely remain elusive. Through assessment of infection dynamics of mallard origin IAV in two different avian hosts, coupled with studies of receptor expression and host response we aimed to reveal the host-pathogen interactions in a cross-species transmission event. We found that shedding patterns and innate immune responses were highly dependent on viral genotypes, host species and inoculation routes, but less dependent on receptor expression. Further, in contrary to the prevailing dogma we demonstrate that birds can produce a wide range of different sialylated structures also found in mammals, e.g. extended *N-* and *O-*linked Neu5Acα2,6 terminated glycans. Overall, receptor incompatibility is not the sole transmission barrier for IAV between birds and to humans, but other host-pathogen factors deserve dedicated studies to achieve proper pandemic preparedness.

---

Zonotic viruses have the capacity to cause severe pandemics ^1^. The three latest influenza A virus (IAV) pandemics (H2N2 1957, H3N2 1968, and H1N1 2009) ^2–4^ have contained genetic material originating from avian influenza viruses. These events have most likely involved spread from wild to domestic bird species and to pigs. Key barriers for interspecies transmission are molecular interactions between host and virus, including recognition of host receptors by virus surface proteins, adaptation to changes in pH and temperature, as well as avoiding the host’s innate and adaptive immune response to infection ^5, 6^.

One of the biological traits required for interspecies transmissions of IAV is a change in receptor binding specificity. Avian viruses preferentially bind to α2,3-linked sialic acid (SA)-containing receptors, whereas human IAVs prefer α2,6-linked SA-containing receptors ^7–11^. Originally it was believed that α2,3-linked SA was exclusive to bird tissues, whereas α2,6-linked SA was exclusive to humans ^12–14^. Furthermore, species specific differences in the sialylated glycan structures serving as receptors for IAVs have been proposed as barriers for transmission between bird species ^12, 13, 15^. More recent characterization of the human airway epithelium has revealed that α2,3-linked SA can indeed be found ^16, 17^, and primary screenings with lectins have indicated the presence of α2,6-linked SA in avian tissues as well ^18–23^. The most important target organs for IAV replication are poorly defined in most bird species. For mallards (*Anas platyrhynchos*), the central reservoir for IAVs, viruses replicate in the surface epithelium of the intestinal tract and transmission occurs via the fecal–oral route^24^. In contrast, the respiratory tract is known to be important for replication in chickens and indirect evidence suggest that this is the case also in an array of wild bird species, including tufted ducks (*Aythya fuligula*) ^23, 25–27^, particularly in the case of infection with highly pathogenic avian influenza virus (HPAIV).

Considering the barriers that dictate cross species transmission events, we hypothesized that the repertoire of expressed sialylated glycan receptors for IAV differs between birds and mammals, and that differences in the glycan structure were to be found even between distantly related bird species. Furthermore, we hypothesized that the outcome of experimental transmission of low pathogenic avian influenza viruses (LPAIVs) between bird species would differ depending on infection route and site of infection, receptor repertoire, host response and virus subtype. Moreover, successful introduction of an LPAIV into a new host species would likely be accompanied by adaptive viral genetic changes. To test these hypotheses, we undertook two main approaches. First, we comprehensively assessed infection potential and host response to infection of a broad array of mallard-origin LPAIVs in two different avian hosts: chicken - a key link to zoonotic transmission, and tufted duck - a representative of the wild reservoir. To evaluate the outcomes of these experiments we assessed virus shedding and monitored the host response as well as SNPs in viruses after infection. Second, we characterized and compared the IAV glycan receptor repertoire displayed on cell surfaces of presumed target organs of mallards, chickens and tufted ducks using glycoproteomics.

## Results

### The outcomes of experimental AIV infection differ between virus subtypes, host species and routes of infection

To interrogate how outcomes of AIV infection can differ depending on host species and route of infection, chickens and tufted ducks were infected with eight different subtypes of mallard-origin LPAIVs (Table S1) using two different routes of virus inoculation: oculonasal or intraesophageal. All birds tested negative for AIV antibodies prior to the infection experiments. All chickens and tufted ducks inoculated intraesophaegally were negative in both oropharyngeal and cloacal swab samples by RT-qPCR for all viruses with only two exceptions: one out of four tufted ducks inoculated with H8N4 showed positive AIV RNA (2.54 10-log EID_50_ equivalent) at 1 day post infection (d.p.i.) and one out of four tufted ducks inoculated with H11N9 (0.79 10-log EID_50_ equivalent) at 2 d.p.i. (data not shown).

In contrast, oculonasal inoculation yielded positive oropharyngeal samples for all viruses in both host species at 1 d.p.i (Figure 1a). In oculonasaly inoculated chickens, virus shedding was detected on all days in all four chickens inoculated with H3N8, H4N6, and H6N2 viruses. In chickens inoculated with H10N1 and H15N5, shedding remained until 3 d.p.i. in a subset of birds whereas the H8N4 virus was detected only at 1 d.p.i. Surprisingly, only one out of four chickens inoculated with the H9N2 virus showed positive AIV RNA (0.58 10-log EID_50_ equivalents), and this was limited to the oropharyngeal sample at 1 d.p.i. Given the high burden of H9N2 in poultry, this is surprising. However, phylogenetically the virus used in this study is located in a distant clade comprising wild bird sequences from Europe, rather than clades dominated by H9N2 viruses in Asian poultry (Figure S1). In chickens, the H6N2 virus showed the most widespread viral replication and AIV RNA was detected in colon in three birds, lungs in two birds, and spleen in a single bird at 3 d.p.i. with virus titre up to 7.27 10-log EID_50_ equivalents. For the other subtypes, viral replication in internal organs were detected to varying extent (Figure S2). In the occulonasally inoculated tufted ducks, all birds shed AIV RNA in oropharyngeal swabs at 1 d.p.i. in all groups except in the H10N1 and H11N9 virus groups, where only two and three birds respectively were positive (Figure 1b). The H6N2 and H9N2 groups showed virus RNA in the oropharyngeal swabs from all tufted ducks until 3 d.p.i. Virus RNA was detected at 3 d.p.i. in the lung (two birds), colon (three birds) and spleen (one bird) of the H6N2 inoculated group (Figure S2). All negative control groups in all experiments demonstrated no signs of disease and no virus was detected in any of the collected swab samples.

**Figure 1.**
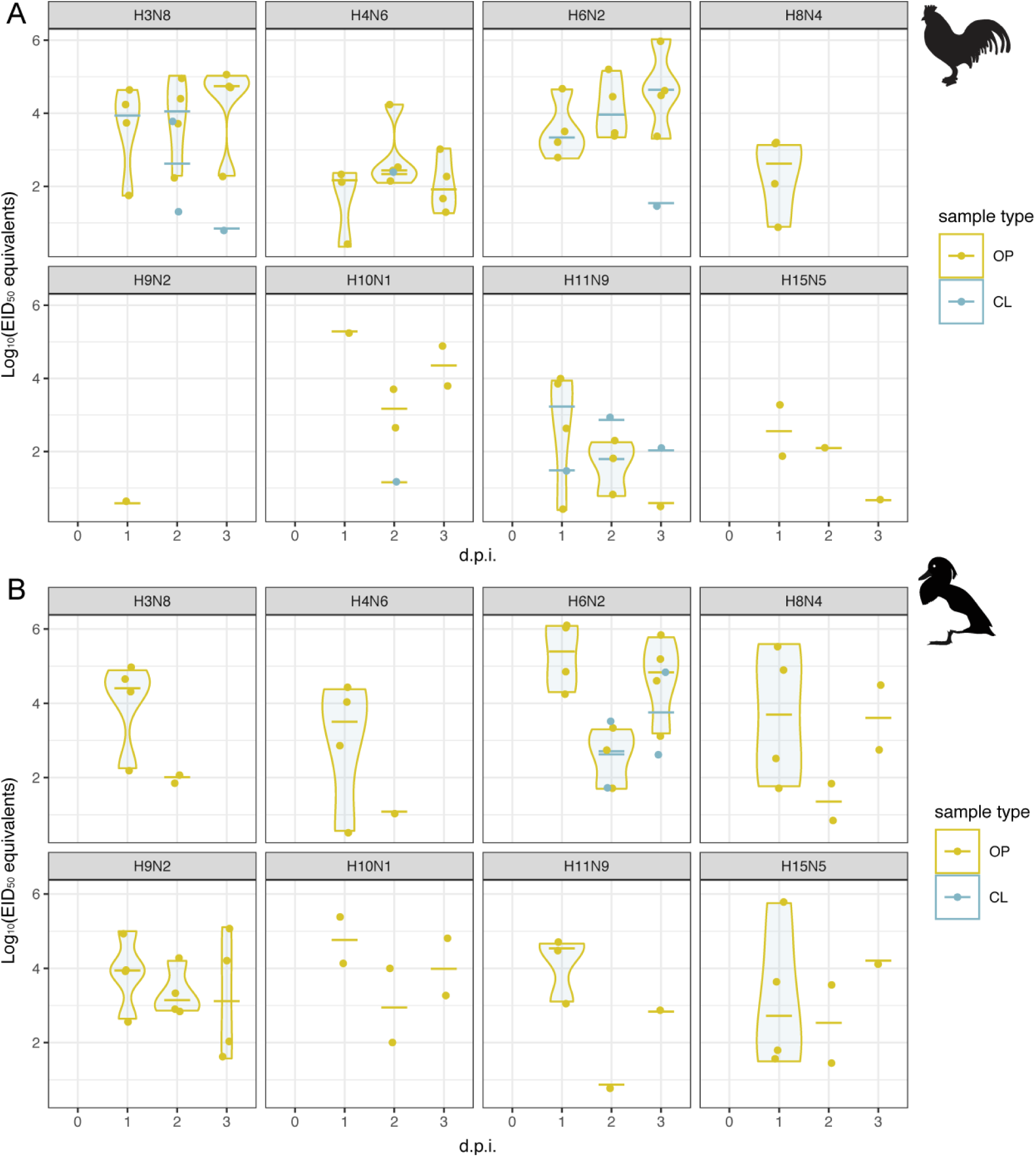
Viral load of mallard origin AIV subtypes in occulonasally inoculated A) chickens and B) tufted ducks. Four birds were inoculated with each virus respectively. Points represent positive samples and horizontal bars their respective median values, and are coloured by sample type; yellow — oropharyngeal (OP) and blue — cloacal (CL) swabs. Violins correspond to kernel density distributions of only positive samples. Negative samples are not shown. Y axis corresponds to log_10_ egg infectious dose 50 equivalents, which are inferred from Cq values based on the relationship reported in Figure S1

Taken together, viral shedding differed depending on AIV subtype, host species and route of inoculation. The intraeosophaegal route produced no virus infections, neither in tufted ducks nor in chickens. This is in contrast to reported results from mallards, where intraeosophaegal inoculation have resulted in cloacal sheding regardless of subtype ^24, 28–30^, and suggests that site of replication might differ between bird species even within the Anatidae family.

### A diversity of single nucleotide polymorphisms in progeny viruses following experimental infection

Putative viral genetic changes during virus infection of chickens and tufted ducks were investigated in swab and tissue samples with virus EID_50_ equivalents high enough to allow for deep sequencing (Table S2). No deletions or insertions were recorded in any of the samples. Previously described amino acid variations associated with host specificity were detected in the NP protein (N319K) of tufted duck/H6 spleen and in the HA protein (N183H) of the tufted duck/H6 cloacal sample at 3 d.p.i. ^31–33^. In addition, a V27L aa substitutional change, previously described in association with amantadine resistance ^34^, was observed in the virus retrieved from tufted duck/H6 colon. Furthermore, SNPs of unkown significance were recovered in the AIV obtained from oropharyngeal swabs from the H3N8 inoculated chicken group as well as the virus recovered from tufted duck/H8 oropharyngeal swab at 3 d.p.i. (Table S2). Finally, the viruses recovered from H9N2 and H10N1 inoculated groups (Table S2) showed no SNPs in their genomes relative to the inoculum.

### Transcriptomics demonstrates differential expression of genes involved in innate immunity and glycosylation in chickens and tufted ducks

In order to understand the host response of chickens and tufted ducks to challenge with mallard-origin LPAIVs, we utilized RNASeq to disentangle gene expression patterns of the hosts. We selected 216 tissue samples comprising lung and colon, as they are important sites of infection, and spleen as it is a primary lymphoid organ. Out of the 216 samples sequenced, 207 were of high quality with an average of 3311801 ± 740763 (mean ± SD) reads per sample. The remaining nine samples were excluded due to low sequencing depth and/or quality.

First, we identified significant differentially expressed genes (DEGs) in each tissue (colon, lung, and spleen) for chickens and tufted ducks for each AIV subtype. In chickens, the number of DEGs ranged from 11–2949 in colon, 34–3225 in lung and 6–3046 in spleen, and in tufted ducks from 49–597 in colon, 7-416 in lung and 92-560 in spleen of birds treated with the different AIV subtypes, see (Figure S3a–b and S4a–b). A marked difference in response to virus challenge in chickens between H3N8 or H4N6 and the other subtypes tested was observed. Fewer DEGs were identified in the tufted duck organs (mean 248 DEGs/organ/virus SD±167), and in contrast to what was observed in chickens, H3N8 was the subtype eliciting the highest number of DEGs. The overlap of DEGs was evaluated between groups infected with the different AIV subtypes within the same species, as well as groups infected with the same subtype between the two species. The proportion of DEGs in response to infection that was unique to each AIV subtype was calculated. In chickens it ranged from 3-32% in colon, 4–24% in lung and 0–31% in spleen across different subtypes (Figure S5a–c). Chickens inoculated with H3N8 had the highest percentage of unique DEGs (32% in colon, 23% in lung, 31% in spleen). The number of unique DEGs detected in a single treatment group in tufted ducks ranged from 6-51% in colon, 0-32% in lung and 10–84% in spleen (Figure S5d–f). As in chickens, H3N8 infected birds displayed the highest percentage of unique DEGs (51% in colon, 31% in lung, 42% in spleen). For all viruses except H3N8 and H4N6, the number of DEGs in response to infection was lower in tufted ducks than in chickens. The overlap of orthologous DEGs between the two species was low to moderate, with H3N8 and H4N6 having a smaller overlap of DEGs between the two species than the remaining virus subtypes (Figure S6-8).

To assess the innate immune response to the LPAIVs used in this study, we specifically studied genes known to be involved in interferone and proinflammatory pathways as well as β-defensins in chickens and ducks at 3 d.p.i ^35^ ^36^. The full list of genes included in this analysis is found in Table S3. In general, up– or down–regulation of such genes was weak or absent in both chickens and tufted ducks. In chickens, a few genes related to interferon signaling were generally affected in response to infection; in colon, *TLR 3* was weakly upregulated for all subtypes except H3N8 and H4N6 (Figure 2, Table S4). However, the adaptor molecule *TICAM1/TRIF* downstream of TLR 3 was generally down–regulated or unaffected in the corresponding colon samples as well as in samples from lung and spleen. The ubiquitine ligase gene, *TRIM25* was weakly up–regulated in response to most IAV subtypes in colon, spleen and lung of chickens. Among the β-defensin genes analyzed, significant changes in response to AIV infection was mainly detected in spleen (and to some extent in lungs) of chickens, where *AvBD1*, *4* and *6* as well as *DEFB4A* were all downregulated at 3 d.p.i. In infected chickens, the H6N2 subtype stood out in evoking the most consistent responses in genes related to IFN signaling (Figure 2, Table S3). From analysis of the tufted duck genome and transcriptome, we could verify for the first time that the RNA sensing protein RIG-I is encoded in this species. However, we could hardly see any transcriptomic response to infection in genes related to IFN signaling or other aspects of innate immunity (data not shown).

**Figure 2.**
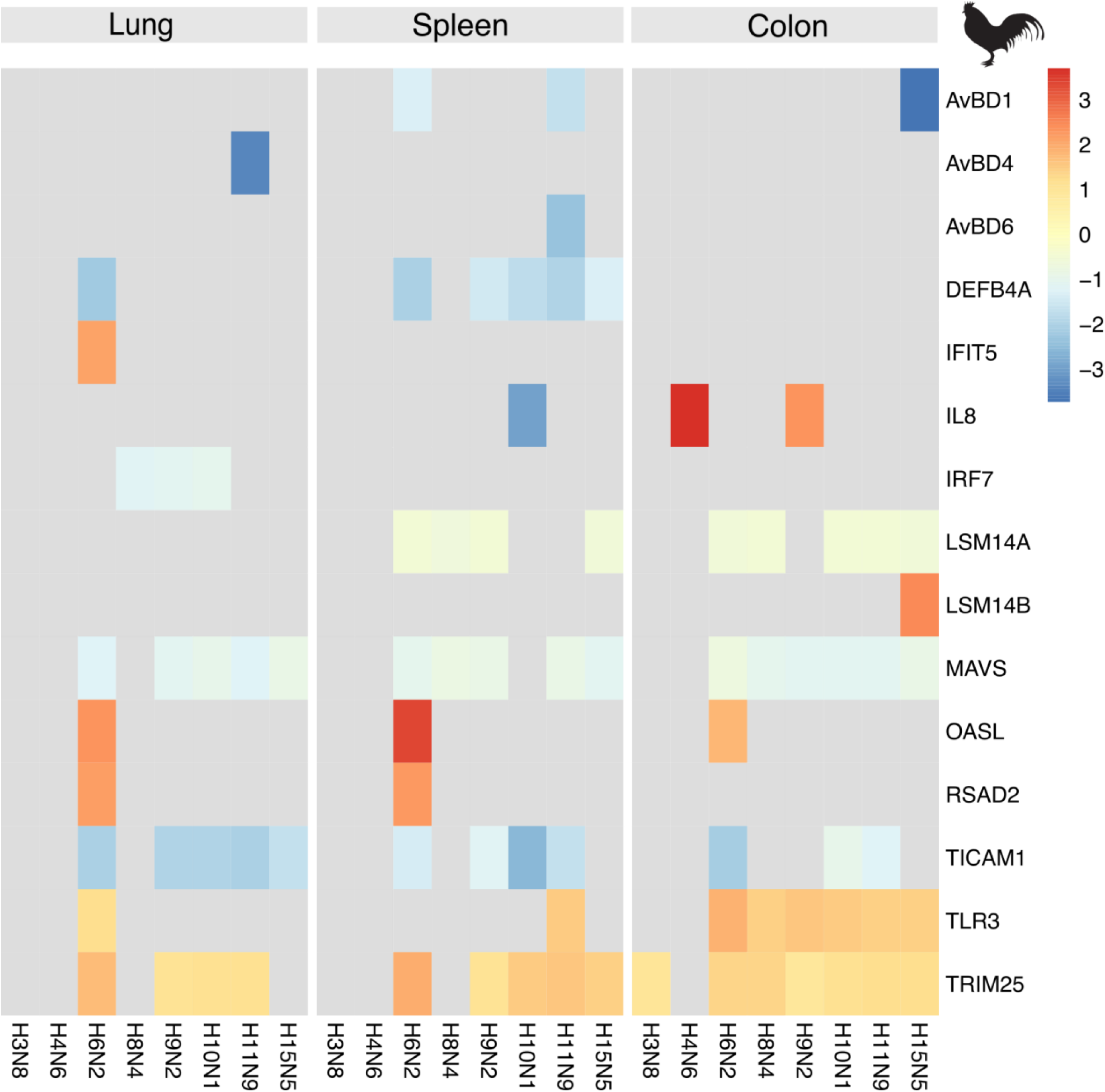
Selected significant DEGs associated with innate immunity in chickens. Genes in the proinflammatory- or RLR signaling pathways as well as β-defensin genes with described functions in birds were selected (Table S1) based on current literature. The color indicates the log_2_ fold change of significant (adjusted p value <0.05 and a threshold of ≥ 10% fold change) DEG’s relative to control birds. Grey indicates non-significant DEGs. Data from lung samples is shown to the left, spleen in the center and colon on the right.

Other genes of note, are the *ANP32* family, serving as co-factors for the virus polymerase during transcription. In chickens, *ANP32E* was weakly down–regulated in all tissues in response to infection with all subtypes except H3N8 and H4N6 (Figure 3a, Table S4). The zinc finger protein ZC3H11A has been suggested to affect replication efficiency of several nuclear replicating RNA viruses including IAV^37^. In our experiments in chickens, *ZC3H11B* was weakly up–regulated for several of the IAV subtypes in all tissues, whereas in tufted ducks *ZC3H11A* was slightly down–regulated in colon for some subtypes (Figure 3a–b, Table S4, S5). Glycosyltransferases related to AIV receptor synthesis were differentially expressed in chickens at 3 d.p.i. Of note was that sialyl transferases adding α2,3-linked SA were generally weakly upregulated in response to infection with most viruses (*ST3GAL2* in spleen, *ST3GAL4* in lung, *ST3GAL5* in spleen) whereas *ST6GAL1* encoding a sialyl transferase adding α2,6-linked SA was weakly down–regulated in all tissues for most viruses. Exceptions to this rule were *ST3GAL6* that was down–regulated in lungs and *ST6GAL2* that was largely unaffected but upregulated in lungs from birds infected with H9N2 and H11N9. A similar pattern could not be observed in tufted ducks (Figure 3b, Table S5). The fucosyl transferase gene *FUT8* was generally up–regulated in all chicken tissues for most viruses whereas *FUT11* was down–regulated, predominantly in the chicken spleen. In tufted ducks, these genes were unaffected except for upregulation of *FUT8* in spleen in response to H4N6 and H8N4.

**Figure 3.**
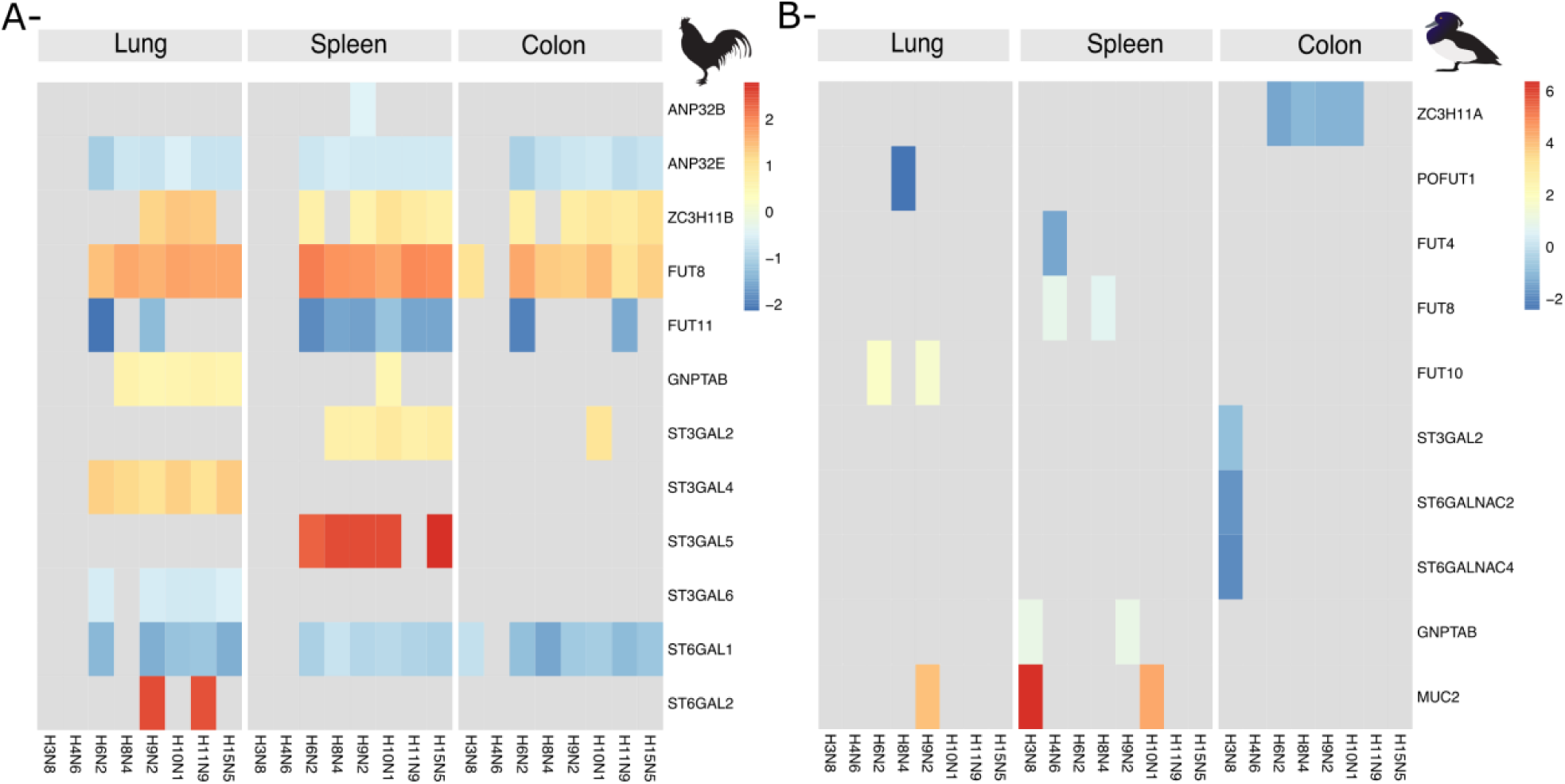
Selected significant DEGs associated with glycosylation and transcription in chickens and tufted ducks. Genes with described association to glycosylation or transcription in birds (Table S2) were selected based on current literature. The color indicates the log_2_ fold change of significant (adjusted p value <0.05 and a threshold of ≥ 10% fold change) DEG’s relative to control birds. Grey indicates non-significant DEGs. Data from chicken (A) and tufted duck (B) lung samples are shown in the left panels, spleen in the center and colon in the right panels.

### Glycoproteomics reveals similarities in influenza virus receptor repertoirs between bird species

To study how the repertoires of sialylated glycan receptors for IAV differ across avian species, we performed glycoproteomic analysis on uninfected specimens of mallards (the original host of the IAV isolates used in this study) as well as of the experimental animals; chickens and tufted ducks. Tissue samples were collected from putative AIV target organs comprising trachea, lung, ileum, and colon from healthy individuals of tufted duck, mallard and chicken. In mallards, *N*-glycopeptides terminating with α2,3-linked Neu5Ac were found in membrane proteins from all sampled tissues (Figure 4 and Table S6). Additionally, *N*-glycopeptides terminating with α2,6-linked Neu5Ac were found in secreted serum proteins from all tissues (Figure 5). However, some *N*-glycopeptides terminating with α2,6-linked Neu5Ac were also detected in predicted membrane proteins. Similar numbers of hits of structures terminated with Neu5Acα2,3 and Neu5Acα2,6 were found in the investigated mallard tissues, except for ileum that had a dominance of Neu5Acα2,3 terminated structures. Disialylated glycoforms carrying one Neu5Acα2,3 and one Neu5Acα2,6 were detected in a smaller fraction of the samples, except for colon that lacked these α2,3/α2,6 mixed sialylated *N*-glycans. The *O*-glycopeptides had core 1 structures (Galβ1,3GalNAcα1–*O*–Ser/Thr) with Neu5Acα2,6 on the GalNAc for serum glycoproteins and Neu5Acα2,3 on the Gal, or both on the Gal and on the GalNAc, for membrane glycoproteins (Figure 4C and 4D). Importantly, longer sialylated *N*-glycans and fucosylated sialylated (S-Le) structures were detected in all sampled tissues from mallards, but concentrated towards ileum (Table S6). In general, there was a dominance of sialylated *N*-glycans compared to *O*-glycans in most tissues except the lungs that contained equal amounts of these structures.

**Figure 4.**
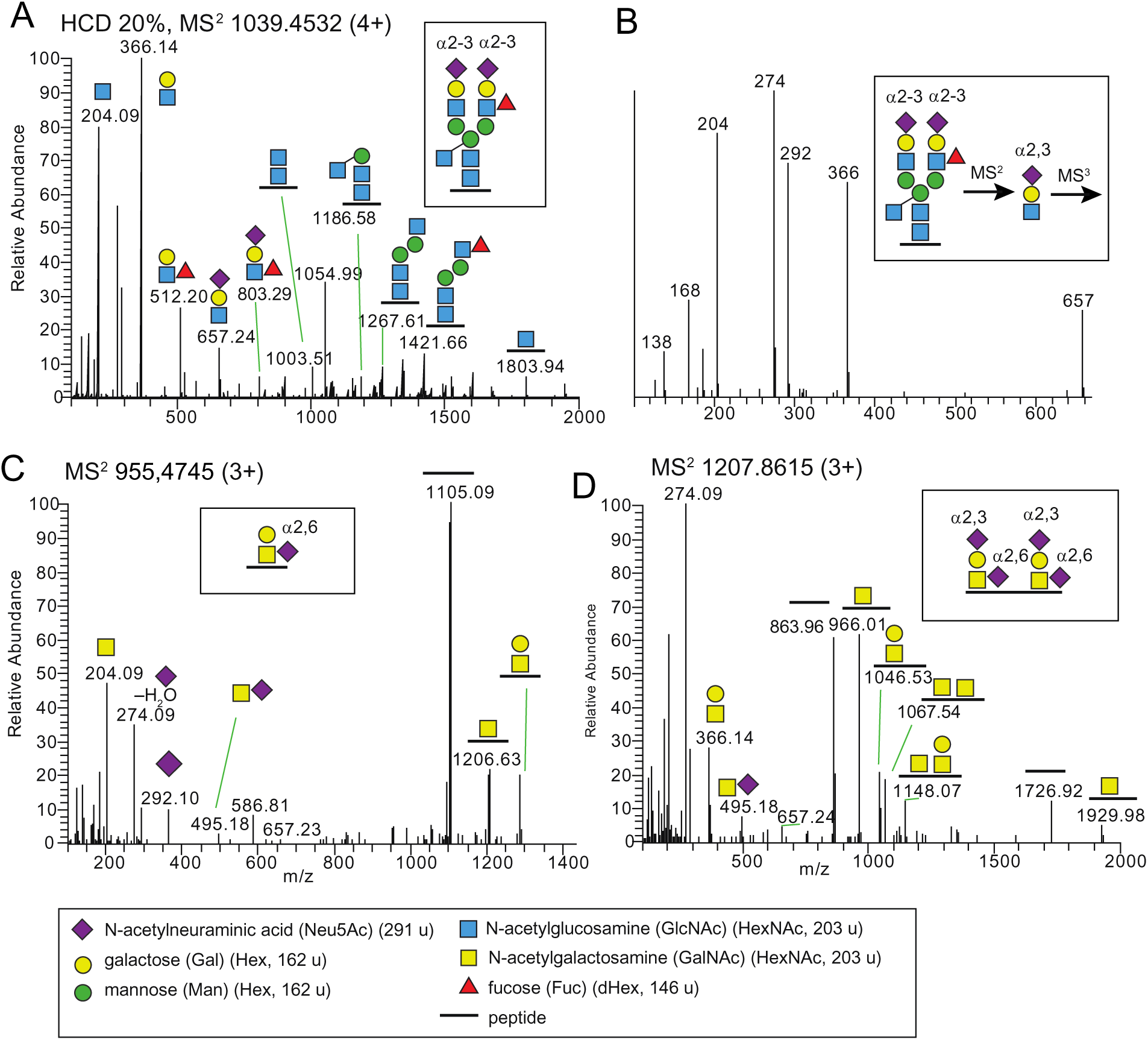
An example of a longer complex disialylated and monofucosylated *N*-glycan and a α2,3/α2,6 core 1 *O*-glycan found in mallards. These glycan epitope structures were also detected in chickens and tufted ducks. (**A**) HCD-MS^2^ of a disialylated complex biantennary N-glycopeptide including a bisecting GlcNAc and a Lewis x fucose (boxed structure) from Epithelial cell adhesion molecule (U3J672_ANAPL) with the sequence R.L<u>N</u>VSIDNEVVQLEK.A. (**B**) MS^3^ of the *m/z* 657 ion provided identification of the Neu5Acα2,3 linkages on both antennas. (**C**) HCD-MS^2^ of a Neu5Acα2,6 sialylated core 1 *O*-glycopeptide from fibrinogen (U3I9E6_ANAPL) with the R.ETAPTLRPVAPPISGTGYQPR.P sequence. (**D**) HCD-MS^2^ of two Neu5Acα2,3 and Neu5Acαα2,6 disialylated core 1 *O*-linked glycans of a glycopeptide from dystroglycan 1 (U3IBQ2_ANAPL) with the R.VISEATPTLAAGKDPEK.S sequence. Glycan symbols are according to the SNFG format ^80^.

**Figure 5.**
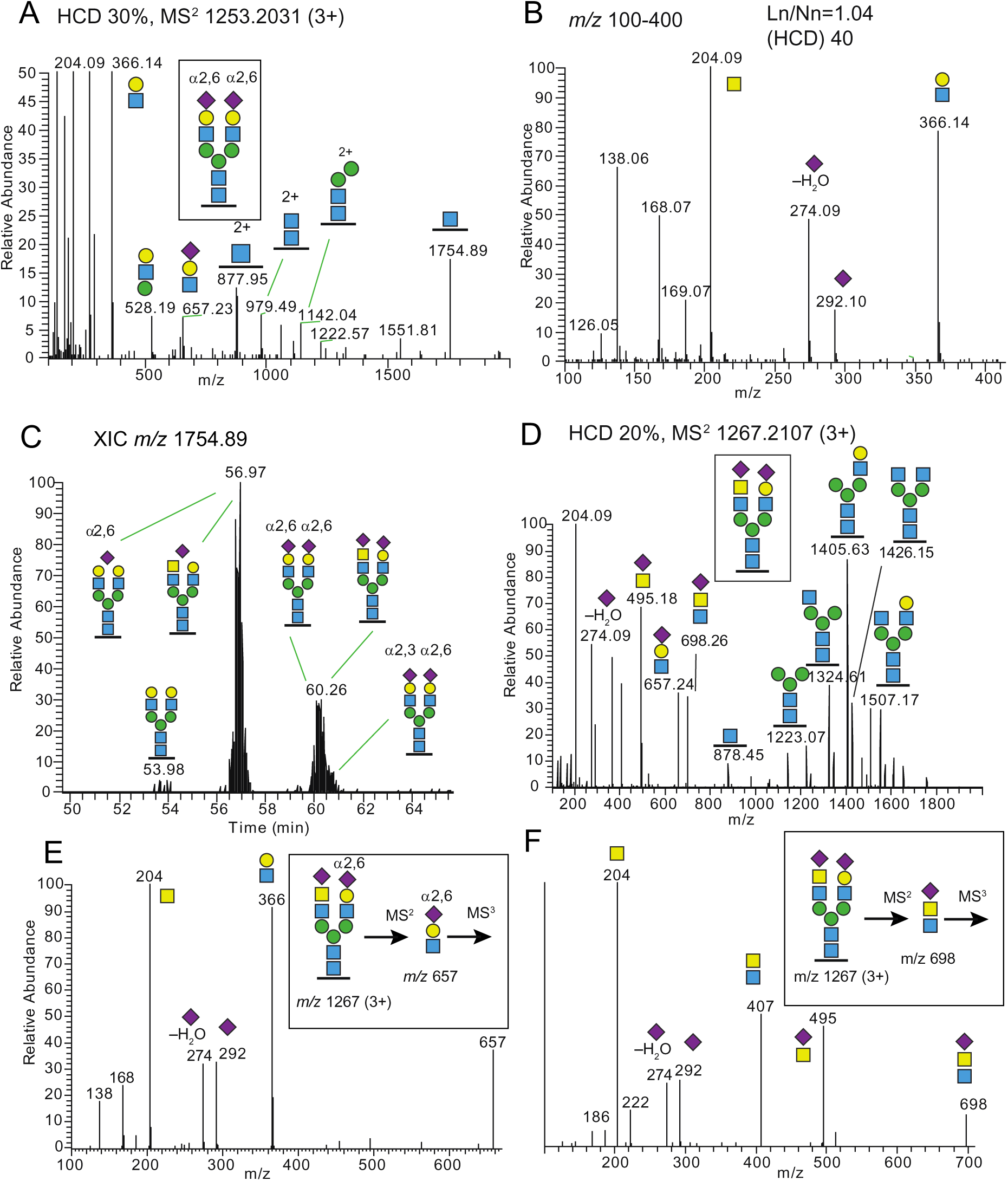
Examples from LC-MS/MS of various Neu5Acα2,6 sialylated *N*-glycopeptides from mallard tissues. These glycan epitope structures were also detected in chickens and tufted ducks. (**A**) HCD-MS^2^ spectrum of a disialylated complex biantennary *N*-glycopeptide (boxed structure) from ovotransferrin (U3J851_ANAPL). (**B**) The *m/z* 100-400 region of the expanded MS^2^ spectrum shows the presence of several oxonium ions used to identify the Neu5Acα2,6 linkage of the sialic acids. (**C**) Extracted ion chromatogram (XIC) of the peptide+HexNAc ion (*m/z* 1754.89) reveals several additional glycoforms of this glycopeptide, e.g. a mixed Neu5Acα2,3; Neu5Acα2,6 glycoform eluting slightly after the major disialylated glycoform at ∼60 min. (**D**) HCD-MS^2^ spectrum of the Neu5Ac_2_Hex_4_HexNAc_5_ glycoform of panel (C) reveals two different antennas. (**E**) MS^3^ of the *m/z* 657.24 ion in panel (D) provides identification of a Neu5Acα2,6 linkage of the HexHexNAc terminated antenna; and (**F**) MS^3^ of the *m/z* 698.26 ion provides identification of Neu5Acα2,6 linkage the HexNAcHexNAc antenna. Glycan symbols are according to the SNFG format ^80^.

In tufted ducks, both transmembrane and secreted glycopeptides terminated with Neu5Acα2,3 or Neu5Acα2,6 were identified (Table S7). Approximately equal ratios of the number of sialylated *N*- vs. *O*-linked glycans were detected in all tissues except for ileum that had dominance of *N*-linked glycans. Fucosylated sialylated glycan structures were detected in all tissues, with the highest amounts in trachea and lowest in ileum. Neu5Acα2,3 was detected in all tissues, with a strong accentuation in trachea (Figure 6a-c). Neu5Acα2,6 was detected in all tissues, but the presence was lower in lung. Disialo glycan structures with both Neu5Acα2,3 and Neu5Acα2,6 were hardly detectable in tufted duck tissues, and only present in trachea and lung at low levels. In chickens, glycopeptides originating from both transmembrane and secreted glycoproteins terminating with Neu5Acα2,3 or Neu5Acα2,6 were identified (Table S8). In trachea, lung, and ileum sialylated *N*-linked glycans dominated and in colon there was a very high dominance of sialylated *N*-linked glycans over sialylated *O*-linked glycans. Fucosylated sialylated glycans were more common in trachea and colon than in lung and ileum. Neu5Acα2,3 was found at similar levels in all sampled tissues, but somewhat concentrated in trachea (Figure 6) and Neu5Acα2,6 was found in similar levels in all sampled tissues.

**Figure 6.**
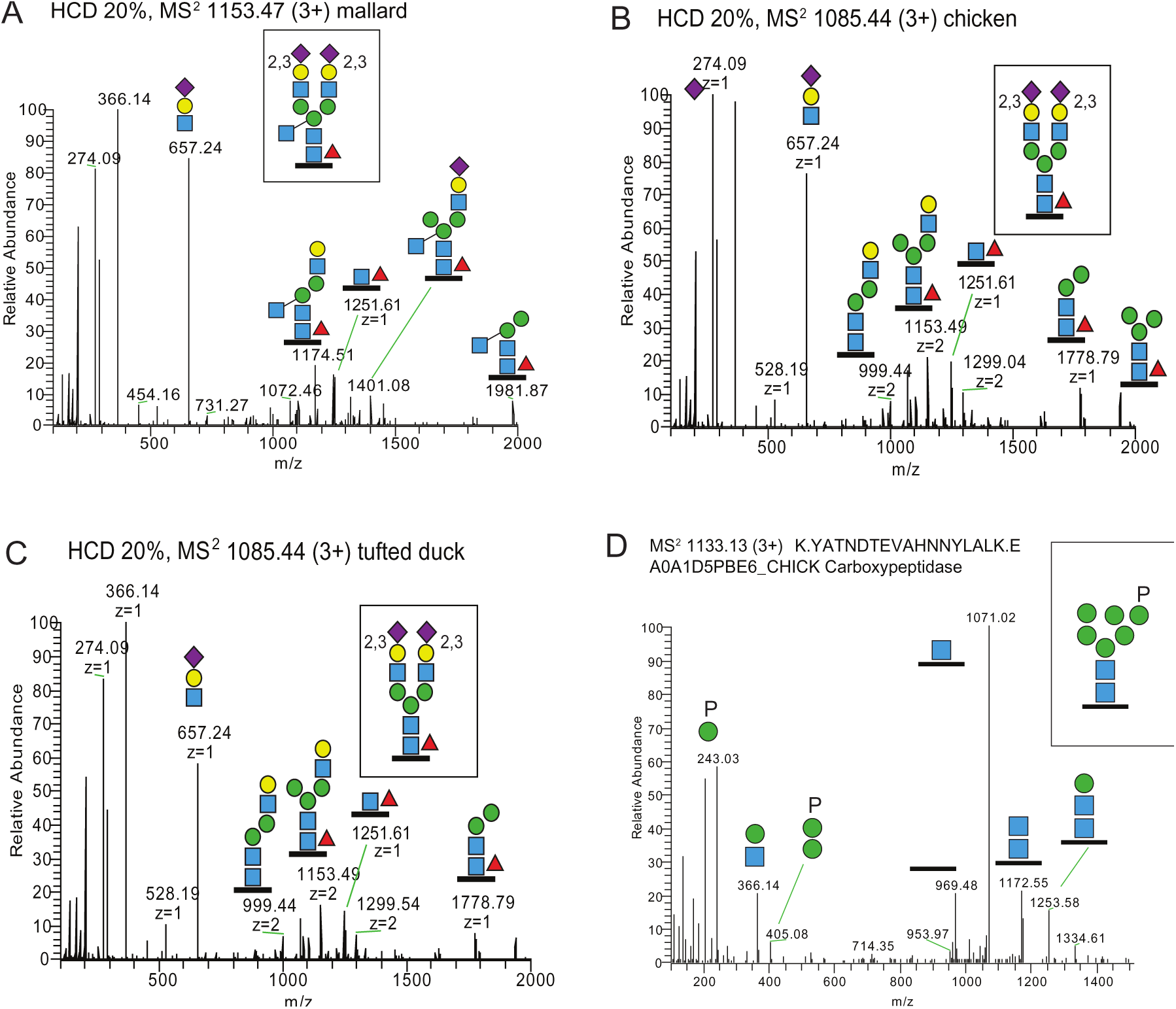
LC-MS/MS of a lumican *N*-glycopeptide and a phosphorylated high mannose glycan detected in bird tissues. The lumican *N*-glycopeptide was found in the trachea of all three species (**A**) mallard, (**B**) chicken and (**C**) tufted duck. The sialic acids are Neu5Acα2,3 linked and the peptide sequence is R.LDGNNLTR.A. (**D**) The phosphorylated high-Man glycopeptide, originated from carboxypeptidase in chicken colon, with the K.YAT<u>N</u>DTEVAHNNYLALK.E sequence. This glycan structure was also detected in mallards (not tested in tufted ducks). Glycan symbols are according to the SNFG format ^80^.

Comparing expressed glycans terminating with α2,3- or α2,6-linked Neu5Ac in different organs between the three species hence revealed great similarities. One important exception was that the abundance of Neu5Acα2,3 terminated fucosylated (sialyl lewis) glycan structures in ileum varied with the highest abundance in mallards and lowest in tufted ducks. In common for all three species, Neu5Gc could not be detected in any of the tissues. Furthermore, despite previously published *in vitro* data suggesting tropism of chicken origin AIV isolates for sulfated sialylated glycopeptides ^12, 15^, no such structures were identified in any of the samples. However, the recently identified putative AIV receptor structures, phosphorylated high-Man *N*-linked glycopeptides ^38, 39^ were identified in both mallard and chicken samples (not tested in tufted ducks) (Figure 6). These glycopeptides were mainly from predicted lysosomal enzymes, including carboxypeptidase and aminopeptidase.

## Discussion

Molecular barriers for transmission of influenza A viruses between birds and to humans can be classified into A) host related factors: e.g. species-specific physiological, biochemical and immunological differences; and B) virus related factors: e.g. subtypes, sublineage specific genetic markers associated with adaptation to a specific host as well as receptor and organ tropism^40^. Mallards are regarded as a key reservoir of AIV, as they carry almost all known AIV subtypes ^41, 42^ and show limited signs of disease even when infected with HPAIV ^43^, which causes significant morbidity and mortality in other bird species, including closely related ducks, such as tufted ducks ^44, 45^, and in chickens ^46^. Through experimental infections together with in-depth molecular characterization of host AIV receptor expression, host response and viral genetic changes during infection, we demonstrate a number of important features central to interspecies transmission.

First, we found that LPAIVs preferentially replicated in the respiratory tract of tufted ducks and chickens. Although this might have been driven by method of inoculation, it is in sharp contrast to LPAIV infection in mallards, where the infection is well-documented in the surface epithelium of the gastrointestinal tract ^24, 47^. The unsuccessful infection of tufted ducks via the intraesophageal route was unexpected as both mallards and tufted ducks belong to the Anatidae family ^48^, and it is long assumed that all duck species share mode of IAV transmission. On the other hand, epizootical studies have shown that AIV prevalence differs dramatically between species within the Anatidae family ^49^. Additionally, studies using virus histochemistry found more virus staining in trachea, rather than colon for several non-mallard species including tufted duck ^23, 25^ suggesting differences in receptor expression. Taken together, our findings suggest that such differences in AIV prevalence between bird species are due to differences in AIV tropism and infectivity, acting as barriers for interspecies transmission.

Second, we found differences in the outcome of infection across the different AIV subtypes tested. Notably, H6N2 was shed at high loads both in oropharyngeal swabs and in tissue samples from both bird species. This virus subtype is very common in wild mallards ^41, 50, 51^ and has been demonstrated to infect gallinaceous birds including pheasants and quails ^52^ as well as chickens ^53^. On the other hand, the unsuccessful infection in chickens with the H9N2 subtype was unexpected as LPAIV H9N2 is endemic in chickens in many countries ^54^. The fact that the HA gene of our H9N2 isolate was phylogenetically distinct from chicken endemic H9N2 isolates suggests the importance of sublineages within the same AIV subtype that seem to be adapted to different host species. Furthermore, in some of the viruses used in this study, we identified putative SNPs that may be related to infection of a new host and provide candidates for futher studies on their role in host adaptation.

Thirdly, transcriptomic analysis illuminated that the host response to virus challenge was different in chickens vs. tufted ducks. For instance, LPAIVs H3N8 and H4N6 generated the least number of DEGs in investigated chickens, whereas the same subtypes yielded the greatest number of DEGs in tufted ducks (see Figure S4). The H6N2 subtype gave the most consistent upregulation of IFN related genes in chickens, consistent with its replication to high titres in lungs, spleens and colons. However, a similar response to this subtype was not detected in tufted ducks despite the presence of the *RIG-I* gene in this species. It should be emphasized though, that the samples in our experiments were taken at 3 d.p.i. whereas in other studies, the strongest innate immune responses to IAV infection in chickens and ducks take place 1-2 d.p.i. ^26, 55, 56^. The up−regulation of *ZC3H11B* in chickens is interesting and merits further study as the A ortholog was described to be involved in nuclear export of mRNA in humans and knocking this gene out in HeLa cells resulted in reduced growth of the IAV strain H1N1 A/WSN/33 ^37^. Taken together, only minor changes in gene expression were identified in virus challenged birds, well in line with earlier reports of LPAIV only inducing mild response in birds ^26, 55, 56^. Particularly, both in terms of number of significant DEGs identified and particular up-/down−regulated DEGs, clear differences could be observed in host response to virus challenge between chickens vs. tufted ducks suggesting a marked difference in host response to virus challenge between species.

Finally, species specific differences in the glycan structures serving as receptors for AIV have been proposed as important barriers for interspecies transmission, most importantly the receptor tropism for α2,3 vs. α2,6-linked Neu5Ac as a hallmark of AIV vs. human adapted IAV ^10, 11, 57^. The results of our glycoproteomic characterization of the avian sialic acid containing glycoconjugates provide a number of important insights into these aspects: Structures containing Neu5Acα2,6 were detected in all sampled avian tissues. In contrast to earlier studies suggesting that avian glycosylation produce shorter, less complex glycans compared to human glycosylation ^58^, we demonstrate that longer sialylated glycan structures were present in all analyzed avian tissues as well (Table S6-9, Figure 6, and Figure S9). Sulfated 3’SLN and S-Le^x^ have been suggested as main receptors for H7 AIVs, H5 AIVs isolated from chickens, and gull-specific H13 and H16 viruses ^12, 13, 59^. However, no sulfated sialylated glycopeptides were identified in any of the investigated avian samples (including chickens), challenging the biological relevance of Sulfated-3’SLN and Sulfated-S-Le^x^ as major AIV receptors. Instead, phosphorylated high-Man *N*-glycopeptides were identified in both the mallard and chicken samples (not tested in tufted duck). This is of high interest as IAVs (of avian, human, and swine origin) recently were described to bind human lung-derived phosphorylated non-sialylated glycans (particularly Man-6P containing glycans) in a SA-independent manner ^38^. We identified these N-glycopeptides on predicted lysosomal enzymes, including carboxypeptidase and aminopeptidase. It can be speculated that IAV binding to phosphorylated glycans takes place during late phase cellular entry, facilitating the virus uncoating process by targeting lysozomal enzymes ^60^. Tropism for *N*-glycolyl-neuraminic acid, Neu5Gc, in wild type AIVs is extremely rare and this structure has not been found in birds ^61^. Indeed, Neu5Gc was not detected in our glycoproteomic analysis in any of the avian species. Taken together, these results not only indicate great similarities in IAV receptor expression between the three bird species, but also demonstrate that avian glycosylation and the expression of glycan structures in avian tissues is more similar to that seen in mammalian tissues than what was previously anticipated.

In conclusion, the experimental design of this study allowed us to address the role of multiple factors as barriers for successful AIV interspecies transmission. The results suggest that not only are factors related to the physiology of the infected species important, but also route of infection, virus phylogeny (as exemplified by the mallard-derived H9N2 LPAIV in chickens) and host response. Moreover, extensive glycoproteomic analysis of the avian respiratory and intestinal tracts clearly demonstrated that birds can produce longer sialylated Lewis structures, as well as both *N*- and *O*-linked glycans terminated with α2,6-linked Neu5Ac. This is contrary to the current dogma whereby these structures were believed to be mammal specific, suggesting that available receptors in mammals and birds are more closely related than previously thought. Receptor incompatibility has been the major focus of AIV interspecies barrier research for the past decades. However, our results suggest that the importance of the proposed receptor barrier might be limited. Instead, we propose that IAV interspecies transmission depends on several host factors (physiology, host immune response, receptor configuration and expression) and viral factors (route of infection, viral phylogeny and genetic makeup). All these factors need to be considered and studied in order to achieve a thorough understanding of IAV interspecies transmission enabling a prudent pandemic preparedness planning.

## Methods

### Ethical statement

All procedures of virus screening and propagation were handled in biosecurity level 2 (BSL2) facilities at the Zoonosis Science Center, Uppsala University. All animal experiments were carried out in strict accordance with a protocol legally approved by the regional board of the animal ethics committee, Sweden (permission number 5.8.18-07998/2017). All animal experiments were conducted in BSL2 animal facilities at the Swedish National Veterinary Institute (SVA).

### Origin of viruses and virus propagation

Viruses used in the current study were obtained from the Linnaeus University AIV repository (Table 1), and collected and isolated as described in ^41^. Briefly, viruses were isolated from mallards captured at a long-term study site at Ottenby bird observatory, Sweden (56°12′ N 16°24′ E). Viruses were propagated in the allantoic cavity of 11-day-old specific pathogen free embryonated chicken eggs and harvested fluid was stored at −70 °C until further use. The egg infectious dose 50 (EID_50_) of the inoculum was determined by infection in embryonated chicken eggs ^62^.

### Strain selection

We selected AIVs comprising eight subtypes of different phylogenetic lineages that have all been detected in the avian reservoir with different frequencies (Table S1). These include those AIVs that are common in mallards at this study site (H4N6 and H6N2) and elsewhere (H3N8), as well as those that are less commonly isolated (H8N4, H9N2, H10N1, H11N9, H15N5). For H15N5 there have only been 14 detections ever defined (https://www.fludb.org/; access 1 July 2020), 6 of which are from this study site. Subtypes seldomly detected in wild bird surveillance (H8N4 and H15N5) may have hosts in undersampled bird species such as diving ducks, although this is still unclear. H9N2 is more common in poultry than in ducks. H10 viruses are isolated from the broadest host range, with infections common in ducks, chickens, seals, and with multiple records of zoonotic infections.

### Pathogenicity and transmissibility in chickens and tufted ducks

A total of 72 white leghorn chickens (*Gallus gallus domesticus*) were raised until 3 weeks old at the SVA. Tufted ducks (*Aythya fuligula*) 3-6 weeks old were obtained from Snavelhof, Veeningen, the Netherlands. All animal experiments were carried out in a biosecurity level two animal facility at the SVA. Birds were divided into groups of four and inoculated with one of the eight viruses or a phosphate buffered saline (PBS) mock. Thirty-six individuals of each species were inoculated intraesophageally (IE) and 36 were inoculated oculonasally (ON), using an inoculum of 10^6^ EID_50_ per 1 mL per bird. For the mock groups, 1 mL of PBS was used per bird. Birds were monitored daily for clinical signs and mortality until 3 days post infection (d.p.i.), when all birds from each group were euthanized by injection of 1 mL of pentobarbital (100 mg). In order to compare virus shedding patterns via the respiratory and/or digestive tract, oropharyngeal (OP) and cloacal (CL) swabs were taken every day from all birds and placed in Virus Transport Media (Glycerol 10% vol/vol; Hanks 1x; Balanced Polymyxin sulphate B 100U/mL; Nystatin 50u/mL; Lactalbumin 5g/L; Penicillin 200U/mL; Streptomycin 200microl/mL; Gentamycin 250microl/mL; MQ water). Tissue samples were collected from lung, colon, and spleen of euthanized birds to study virus distribution as well as the transcriptomic host response to infection.

### Reverse Transcriptase quantitative real-time-PCR

Viral RNA was extracted from the OP and CL swab samples collected from chickens and tufted ducks using Maxwell® 16 Viral Total Nucleic Acid Purification Kit on a Maxwell® 16 System extraction robot (Promega, Madison, WI, USA). RNA was isolated from lung, colon and spleen of all ON inoculated birds using TissueLyser II and RNeasy Mini Kit (Qiagen, Hilden, Germany). Quantification of the viral load (expressed as egg infective dose 50 (EID_50_) equivalent, as defiend below) from the swab samples was based on Cq values obtained by reverse transcriptase quantitative real-time PCR (RT-qPCR) targeting the matrix (M) gene ^63^ and using AgPath-ID One-Step RT-PCR reagent kit (Thermo Fisher Scientific, Waltham, MA,USA) on a a BioRad CFX1000 Real-Time PCR System (BioRad, Hercules, CA, USA).

We calculated EID_50_ equivalent by determining the Cq values from a known EID_50_ reference virus inoculum. From this data we generated a standard curve (Figure S10) correlating these measurements and used this curve to predict EID_50_ equivalents from Cq data for all virus experiments.

### Whole genome sequencing and variant detection

Total viral RNA was extracted from the virus inocula, all organs with a Cq value < 30, and positive swabs at 3 days post infection (d.p.i.) with a Cq value < 30 (Table 1) using QIAamp viral RNA Mini kit (Qiagen, Hilden, Germany) according to the kit protocol. Virus whole-genome amplification was undertaken as described by Pohlmann, et al. ^64^. Sample quality control was assessed using a Qubit 4 Fluorometer (using Qubit™ dsDNA BR assay Kit) and Agilent 2200 TapeStation (using Agilent D5000 ScreenTape assay kit). Library construction and sequencing was undertaken at Novogene (Beijing, China). The genomic DNA was randomly fragmented to size of 350bp, and then DNA fragments were end polished, A-tailed, and ligated with the adapters of Illumina sequencing, and further PCR enriched with primers of P5 and P7 oligos. The PCR products as final construct of the libraries were purified followed by quality control test. 150 bp paired reads were generated with a sequencing depth of 9-12 million reads per sample on the Illumina platform NovaSeq 6000.

For variant detection, generated reads were analysed using Geneious Prime work package (Biomatters, Auckland, New Zealand) through the following pipeline: Initially, the primer sequences were trimmed off from the raw reads using the “Trim Ends” Geneious Prime plugin. Further, the trimmed reads were mapped using bowtie2 implemented in Geneious Prime against the whole genome sequence obtained from the used homologous inoculum, and a consensus sequence was generated for each sample. Finally, SNP (single nucleotide polymorphism) calling was conducted using “variation/SNPs” plugin implemented in Geneious Prime work package on the assembled contigs with a maximum variant p-value of 10^6.

### Serology

To confirm that all birds were seronegative prior to infection, blood samples were collected using vacutainer serum plus blood collection tubes (BD, Becton, Dickinson and Company, New Jersey, USA) from all animals 2 days prior to the first animal experiment. Serum was recovered by centrifugation of the tubes at 1000 g for 5 minutes. Samples were examined for AIV-specific antibodies using the Influenza A nucleoprotein (NP) antibody competition ELISA kit (Idexx, Hoofddorp, The Netherlands) according to the manufacturer’s instructions.

### Transcriptomics

Gene expression was quantified in lung, colon and spleen for all birds three d.p.i using QuantSeq 3’ mRNA sequencing ^65^. For this purpose, 25 mg of each tissue was homogenized using Qiagen TissueLyser II, and RNA was extracted using RNeasy Mini Kit and DNAse treated using the RNase-Free DNase Set (Qiagen, Hilden, Germany). The quantity and quality of the RNA was assessed using the Qubit (Qubit RNA BR Assay Kit, Invitrogen, CA, USA) and the Agilent 2100 Bioanalyzer system (Agilent RNA 6000 Nano Kit, CA, USA). The library preparation was conducted using the QuantSeq 3‘ mRNA-Seq Library Prep Kit (FWD) (Lexogen Inc., Greenland, USA) according to the manufacturer’s instructions. Samples were sequenced (75bp single-end reads) on the Hiseq 4000 platform (Illumina Inc., CA, USA). The raw QuantSeq reads were trimmed using the BBDuk program in the package BBMap (https://sourceforge.net/projects/bbmap/) to remove adapter sequences, poly-A tails, overrepresented sequences, and low-quality bases with the following parameters: k = 13, ktrim = r, useshortkmers = t, mink = 5, qtrim = r, trimq = 10, minlength = 20. Read quality before and after trimming was checked using FASTQC 0.11.5 ^66^. The trimmed sequences were aligned to the chicken reference genome (GRCg6a, Ensembl release 99, accession number GCA_000002315.5) and the tufted duck reference genome (bAytFul2.pri, GenBank release 236, accession number GCA_009819795.1) using STAR 2.5.3a ^67^. For the STAR indexing step, the tufted duck gff annotation file from NCBI was converted into gtf format using the gffread utility in the Cufflinks software 2.2.1 ^68^. Read counts per gene were calculated in STAR 2.5.3a.

Bioconductor DESeq2 v1.24.0 ^69^ program in R v3.6.1 (RCoreTeam, 2016) was used to calculate statistical differences of the expression levels of genes between the control birds and the infected birds for each species and tissue (four individuals per group). Cut-off values for significant genes were set to an FDR of <0.05 and to genes that were up- or downregulated more than 10%, based on visual inspection of volcanoplots generated in the EnhancedVolcano package ^70^ in R v3.6.1 ^71^.

To allow for comparisons of gene expression levels between chicken and tufted duck, a best hit (RBH) analysis was conducted. For this purpose, the DNA sequence information for coding sequences (cds) from chicken (GRCg6a, Ensembl release 99) was downloaded. The gffread utility from the STRINGTIE package ^72^ was then used to generate a FASTA file with the DNA sequences for the coding sequences in the tufted duck. The coding sequences were reciprocally blasted using blastn ^73^. RBHs were identified using the python script reciprocal_blast_hits.py https://scriptomika.wordpress.com/2014/01/28/extract-best-reciprocal-blast-matches/.

To compare the response between treatment groups the overlap of differentially expressed genes between groups treated with different virus subtypes within one species, and between groups treated with the same virus subtypes between the two species were visualised using the UpSetR package v 1.4.0 ^74^ in R v3.6.1 ^75^.

### Transcriptomic response and gene onthology

Gene expression levels for genes associated with innate immune response, transcription, glycosylation, and inflammation were visualized in heatmaps. Significantly differentially expressed genes associated with these functions were identified by filtering the DESeq2 output files for gene names identical to the list disclosed in supplementary Table S1 or gene names including any of the search terms disclosed in supplementary Table S2 and S3 or gene descriptions including “defensin” or “gallinacin” (for β-defensins/gallinacins). A threshold for significant DEGs was set at ≥ 10% fold change relative to control birds and an adjusted p-value ≤ 0.05. Gene ontology and pathway analysis were undertaken for differentially expressed genes (adjusted p-value ≤0.05) for each virus/bird species/organ. Gene set analysis/overrepresentation analysis was conducted using the Webgestalt web interface (http://webgestalt.org/) searching the gene ontology database for “molecular function”, “cellular component”, and “biological process”. Additionally, pathway analysis was conducted searching KEGG using the Webgestalt web interface as well. Ontology terms/pathways with a FDR≤0.05 were visualised in heatmaps.

### Epithelial cell isolation for glycoproteomic analysis

Tissue samples were collected from trachea, lung, ileum, and colon of captive healthy uninfected mallards, chickens and tufted ducks (5 individuals per species). The biopsies were rinsed in chilled (∼8 °C) PBS (Sigma-Aldrich AB, Stockholm, Sweden) upon collection to remove any debris. Adipose tissue, blood vessels, etc. was removed and the intestinal specimens were cut open and intestinal content and mucus was carefully removed. Cleaned tissue samples were incubated at 37 °C for 1 h in 10 mL DPBS with 1 mM DTT and 3 mM EDTA (Sigma-Aldrich). After incubation, the liquid was carefully aspirated and replaced with PBS (Sigma-Aldrich). The samples were vortexed for 5×20 s and the tissue was then removed. The remaining cell slurry was spun at 1000 g for 5 min at 4 °C and the supernatant was discarded. The pellets were snap-frozen in liquid nitrogen and stored at −80 °C.

### Protein digestion, enrichment and fractionation

The samples were homogenized using the lysis matrix D on FastPrep®-24 instrument (MP Biomedicals, OH) in lysis buffer (50 mM triethylammonium bicarbonate (TEAB), 2% sodium dodecyl sulfate (SDS)) and 5 cycles 40 s each. The samples were centrifuged at maximum speed for 15 min, and supernatants harvested. The protein concentration was determined using Pierce™ BCA Protein Assay (Thermo Fisher Scientific) and the Benchmark Plus microplate reader (BIO-RAD) with BSA solutions as standards.

Sample aliquots (500–1000 µg) were digested with trypsin using the filter-aided sample preparation (FASP) method ^76^, with small modifications. Briefly, samples were reduced with 100 mM dithiothreitol at 60°C for 30 min, spin-filtered (30 kDa MWCO Pall Nanosep centrifugation filters, Sigma-Aldrich), washed repeatedly with 8 M urea and followed by digestion buffer (1% sodium deoxycholate (SDC) in 50 mM TEAB) prior to alkylation with 10 mM methyl methanethiosulfonate in digestion buffer for 20 min. Protein digestion was performed in digestion buffer by addition of 5 µg Pierce MS grade Trypsin (Thermo Fisher Scientific) at 37°C overnight, followed by an additional 2 h incubation with new trypsin added the consecutive day. Peptides were isolated by centrifugation and SDC was removed by acidification with 10% trifluoroacetic acid. Glycopeptides were enriched with hydrophilic interaction liquid chromatography (HILIC) according to ^77^, with slight modifications. In short, peptides were loaded onto an in-house zwitterionic Zic-HILIC SPE cartridge containing 20 mg of Zic-HILIC particles (10 μm, 200 Å; Sequant/Merck). The flow-through was collected and re-circulated through the column an additional three times. The column was washed with totally 1.2 mL of 80 % (v/v) acetonitrile and 1 % (v/v) trifluoroacetic acid. Enriched glycopeptides were eluted with 4 times 50 µL 0.1 % (v/v) trifluoroacetic acid followed by 50 μL of 25 mM NH_4_HCO_3_ and finally 50 μL of 50 % (v/v) acetonitrile and dried by vacuum centrifugation. The samples were fractionated into six fractions (5.0-17.5 % acetonitrile in 0.1% trimethylamine), using Pierce high pH reversed-phase peptide fractionation kit (Thermo Fisher Scientific) according to the manufacturer’s protocol and pooled into 3 samples. Pooled samples were dried in a vacuum centrifuge and reconstituted in 15 μL of 3 % acetonitrile, 0.1 % formic acid for LC-MS/MS analysis.

### NanoLC MS analysis

Peptide samples were analyzed on an Orbitrap Fusion Tribrid mass spectrometer interfaced with Easy-nLC1200 liquid chromatography system (Thermo Fisher Scientific). Peptides were trapped on an Acclaim Pepmap 100 C18 trap column (100 μm x 2 cm, particle size 5 μm, Thermo Fischer Scientific) and separated on an in-house packed analytical column (75 μm x 30 cm, particle size 3 μm, Reprosil-Pur C18, Dr. Maisch) using a linear gradient (solvent A; 0.2 % formic acid in water and solvent B; 80 % acetonitrile, 0.2 % formic acid in water) from 7 % to 35 % of solvent B over 45 min followed by an increase to 100% solvent B in 5 min, and finally 100% solvent B for 10 min at a flow rate of 300 nL/min. MS^1^ scans were performed at 120 000 resolution, *m/z* range 600-2000, the most abundant double or multiply charged precursors from the MS^1^ scans were selected with a duty cycle of 3 s, isolated with a 3 Da window, fragmented with higher-energy collision induced dissociation (HCD) at 30% normalized collision energy (NCE), *m/z* range 100-2000, and then two times at 40% NCE, *m/z* range 100-2000 and *m/z* 300-2000. The maximum injection time was 118 ms and MS^2^ spectra were detected in the Orbitrap at 30 000 resolution. Dynamic exclusion was enabled with 10 ppm tolerance and 10 s duration. For selected samples, MS^3^ was performed on the *m/z* 657.23 ion [Neu5AcHexHexNAc]^+^ using HCD 20% NCE at both the MS^2^ and the MS^3^ steps, and with detection in the ion trap.

### Glycoproteomic data analysis

The LC-MS/MS raw files were analyzed with the Byonic software (Protein metrics) using a modified list of the glycan modifications “182 human N-glycans” and “6 most common O-glycans”, such that e.g. glycoforms with a Hex_4_HexNAc_5_ core structure, differing from the normal Hex_5_HexNAc_4_ complex biantennary core, were added. Further Byonic search criteria: the FASTA database was *Anas platyrhynchos* (Organism ID 8840; 27089 sequences) and *Gallus gallus* (Organism ID 9031); C-terminal cleavage allowed after Lys and Arg; accuracy for MS^1^ was 10 ppm and for MS^2^ it was 20 ppm; static modification was methylthio on Cys (+45.9877 u) and variable modification, apart from glycans, was Met oxidation (+15.9949 u). Tufted duck glycopeptides were identified using the mallard genome as template, due to non-available tufted duck genome at the time of analysis. A Byonic score cut off >300 was used for glycopeptide hits and Neu5Ac containing hits were manually verified with the following inclusion criteria 1) presence of the correct peptide+HexNAc ion with respect to the precursor mass and identified glycan mass; 2) presence of the Neu5Ac oxonium ions *m/z* 274 and *m/z* 292; and 3) for hits including fucose (dHex), presence of peptide+HexNAc+dHex ion for the identification of a core fucose, and/or presence of *m/z* 512 ion (HexHexNAcdHex) for identification of Fuc on the antennae. Extracted ion chromatograms were traced at diagnostic MS^2^ ions including peptide+HexNAc ions, chosen to identify additional glycoforms sharing the same peptide, and saccharide oxonium ions, for instance *m/z* 274.09 for tracing Neu5Ac; *m/z* 290.09 for tracing Neu5Gc and *m/z* 495.18 for tracing Neu5AcHexNAc.

The Neu5Acα2,3/Neu5Acα2,6 isomeric structures of glycopeptide hits were determined according to ^78^.. Briefly, the relative LacNAc to Neu5Ac (L/N) oxonium ion ratio (*m/z* 204 + *m/z* 366) / (*m/z* 274 + *m/z* 292) were calculated at an NCE of 40% for *N*-glycopeptide hits carrying Hex5HexNAc4NeuAc_2_(dHex)_0-1_ complex biantennary structures; and for *O*-glycopeptide hits carrying HexHexNAc(Neu5Ac)_1-2_ structures. The ratio was then multiplied with the stoichiometric compensation factor *n*(NeuAc) / *n*(HexNAc) to obtain the normalized ratio Ln/Nn. For the MS^3^ based methodology, the MS^3^ spectrum of the MS^2^ generated *m/z* 657.23 ion was manually inspected. When the (*m/z* 204 + *m/z* 366) ion intensities were larger than the (*m/z* 274 + *m/z* 292) intensities, the linkage structure was considered as Neu5Acα2,6 and when smaller the linkage structure was considered as Neu5Acα2,3. For the MS^3^ experiments, Byonic hits with scores <300 were also considered, provided that the same hit was present with a score >300 in the corresponding MS^2^ experiment.

## Data and Code Availability

The authors declare that all data supporting the findings of this study are available within the article, its supplementary information files, or are available from the authors upon request. The nucleotide sequences generated for variant detection and transcriptomic analyses (RNAseq) reported in this paper have been deposited into public databases, Sequence Read Archives, under project number PRJNA664709.

The mass spectrometry proteomics data have been deposited to the ProteomeXchange Consortium via the PRIDE ^79^ partner repository with the dataset identifier PXD021392.

## Acknowledgments

We acknowledge Josanne Verhagen for culturing viruses and Janina Krambrich for her help in the preparation of samples for NGS analysis. We acknowledge Dr. Roy Francis (NBIS, Science for Life Laboratory) for helpful input on bioinformatics. The glycoproteomic analysis was performed at the BioMS node at the Proteomics Core Facility of Sahlgrenska Academy, University of Gothenburg. Support from the Swedish National Infrastructure for Biological Mass Spectrometry (BioMS), funded by the Swedish Research Council (VR), is gratefully acknowledged. We are grateful of Inga-Britt and Arne Lundbergs Forskningsstiftelse for the donation of the Orbitrap Fusion Tribrid MS instrument. The transcriptomic analyses were performed using computational resources at the bwUniCluster funded by the Ministry of Science, Research and the Arts Baden-Württemberg and the Universities of the State of Baden-Württemberg, Germany, within the framework program bwHPC-C5.

## Author contributions

Conceptualization: PEl, JJ, BO, AL, MW, MN, PEr

Animal experiment: MN, PEr, PEl, JJ, CL, CB

Transcriptomic analysis: EJ, PEr, RHSK, MN

Glycoproteomic analysis: JN, PEr, CS, BMO, GL

Writing — original draft: MN, PEr, PEl, EJ

Writing — review & editing: MN, PEr, PEl, JJ, AL, BO, JN, GL, MW, EJ, CL

## Supplementary Materials

### Supplementary figures

**Figure S1.**
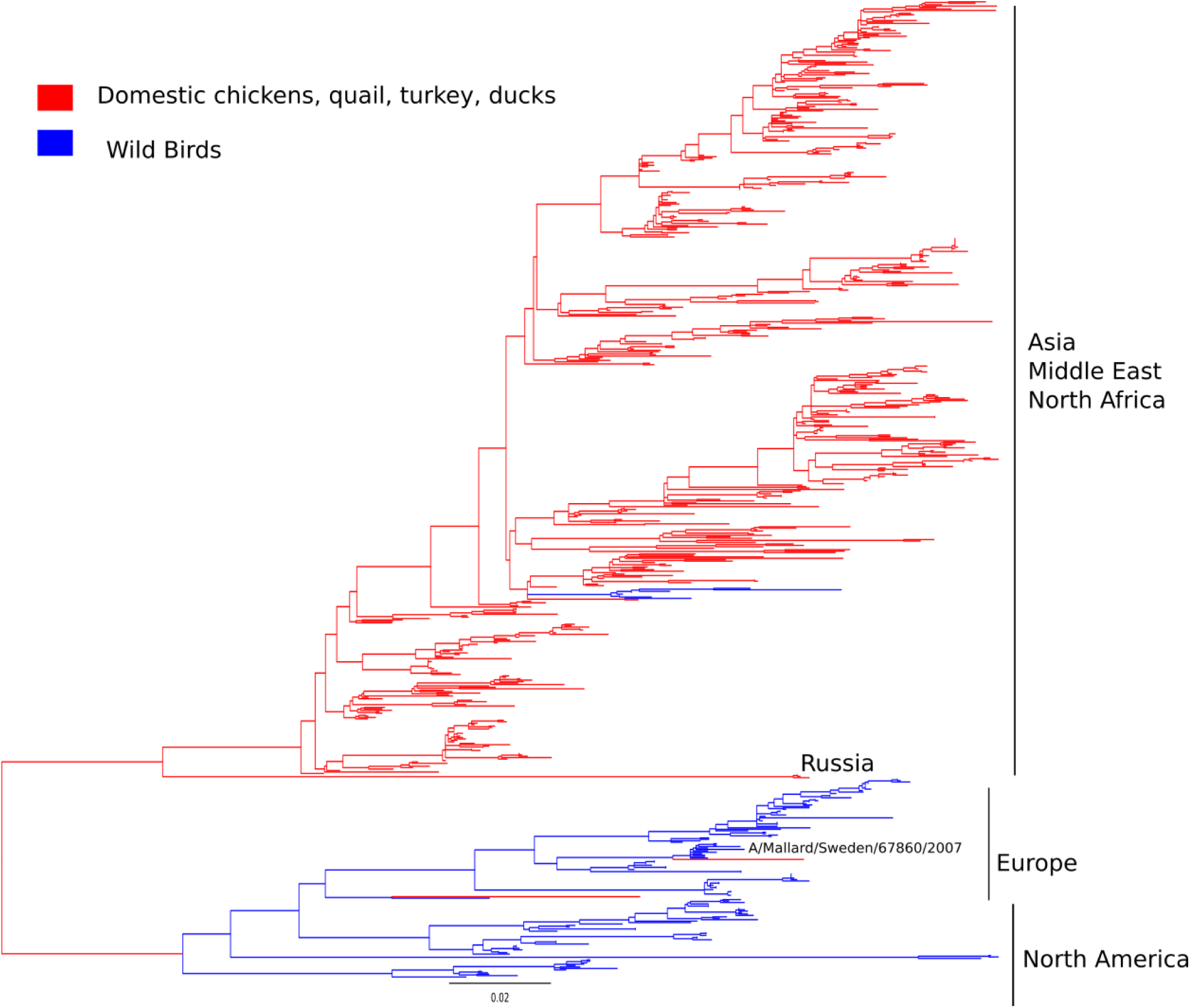
LPAIV H9 Phylogeny. Maximum likelihood tree of the HA gene segment of influenza A(H9N2) virus. Sequences obtained from domestic chickens, quail, turkeys, and ducks are coloured in red. Sequences obtained from wild birds are coloured in blue. Geographic location is provided to the right of the tree. Scale bar indicates number of nucleotide substitutions per site. Tree is midpoint rooted, reflecting the long branch length between the poultry-associated clade and those viruses from Europe and North America.

**Figure S2.**
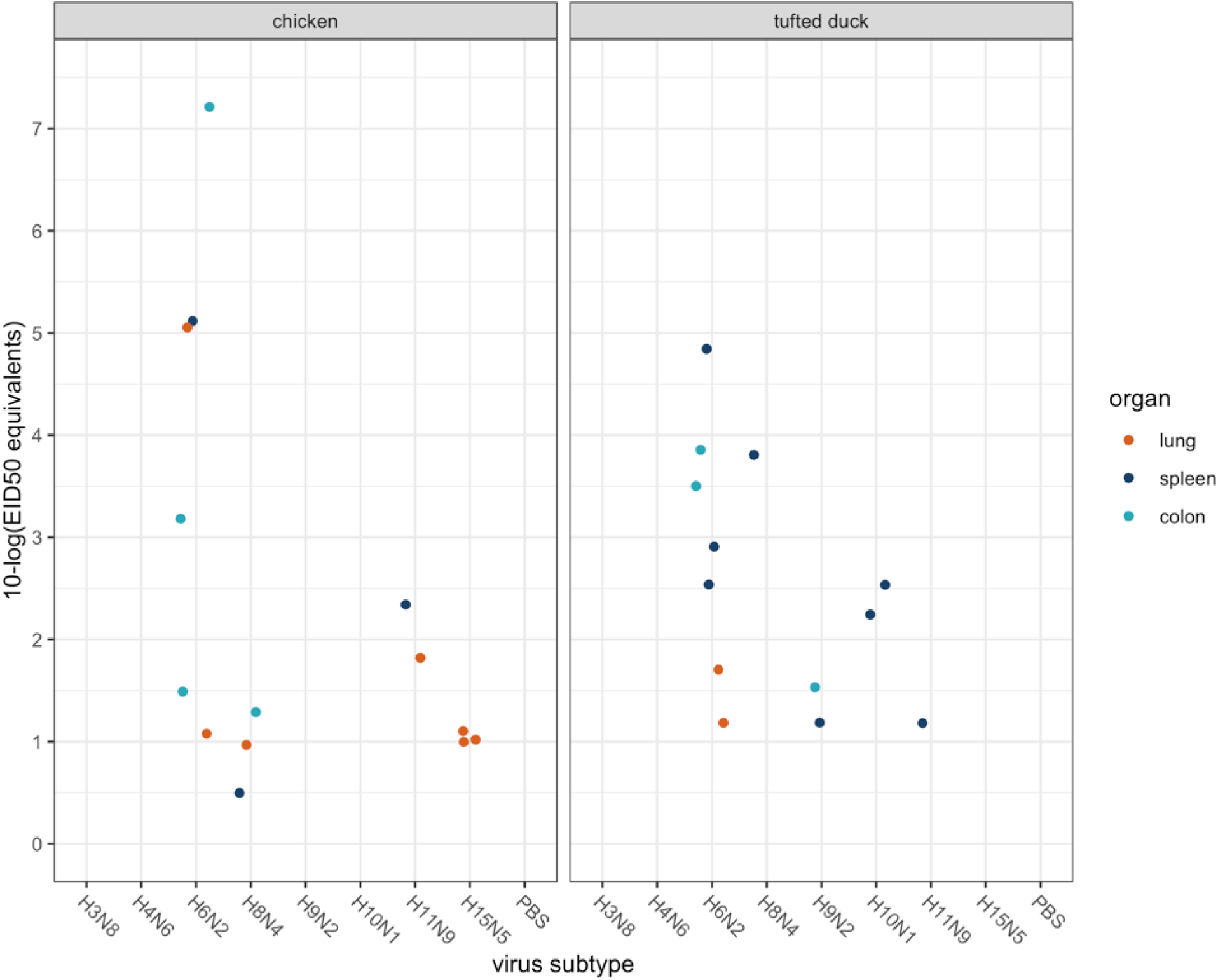
Strain specific AIV positive organ samples. Scatter plot of obtained positive organ samples at 3 days post infection from ON inoculated chickens and tufted ducks. Each point illustrates one positive sample from one individual. Four individuals were inoculated with each virus respectively. The points are coloured based on organ type; orange — lung, dark blue — spleen, and light blue — colon. Log_10_ egg infectious dose 50 equivalents vs. inoculum.

**Figure S3.**
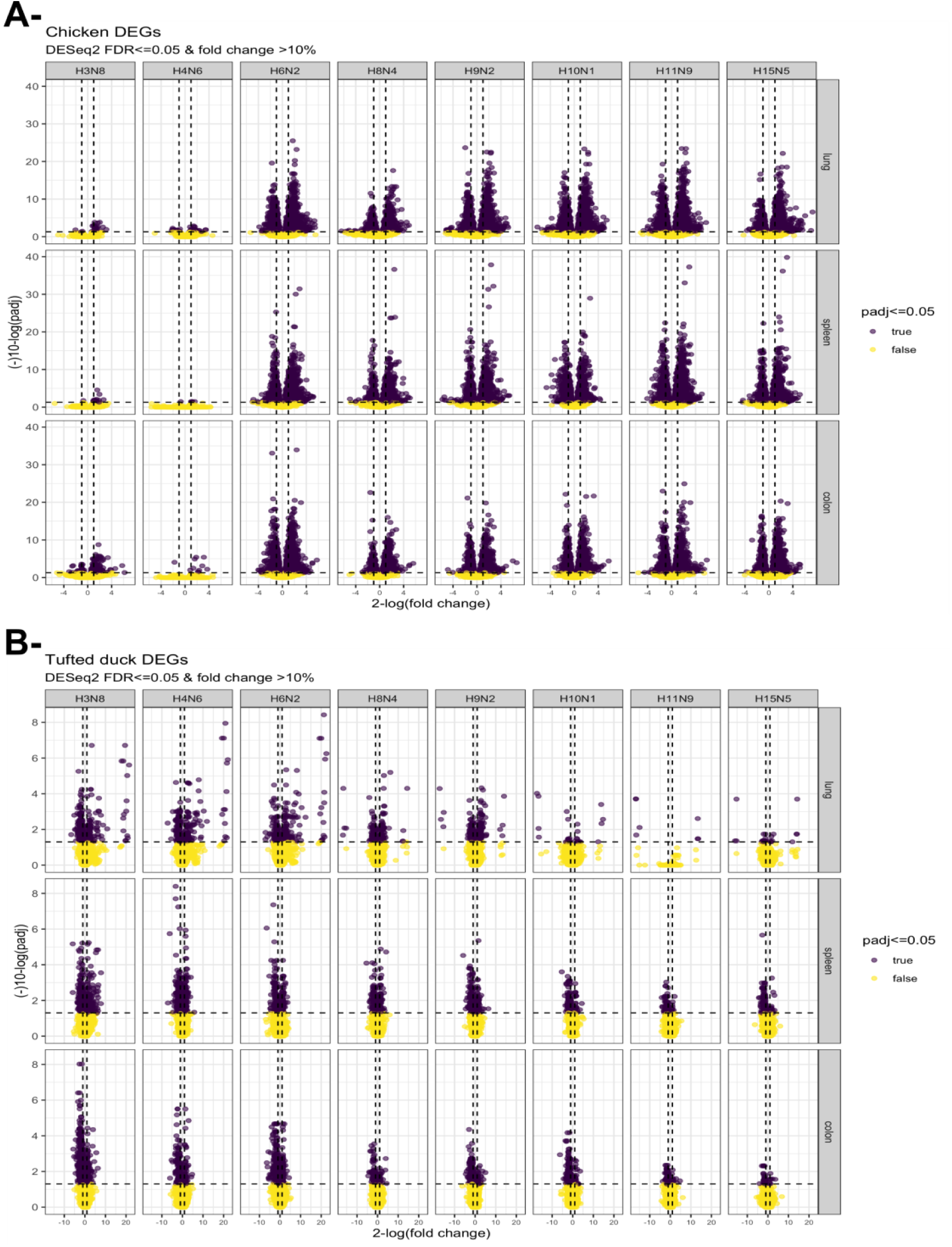
Volcano plots obtained from chickens (a) and tufted ducks (b) infection experiments. Volcano plot showing DEGs for H3N8, H4N6, H6N2, H8N4, H9N2, H10N1, H11N9, and H15N5. The x-axis represents the log2 values of the fold change observed for each mRNA transcript, and the y-axis represents the log10 values of the adjusted p-values <=0.05.

**Figure S4.**
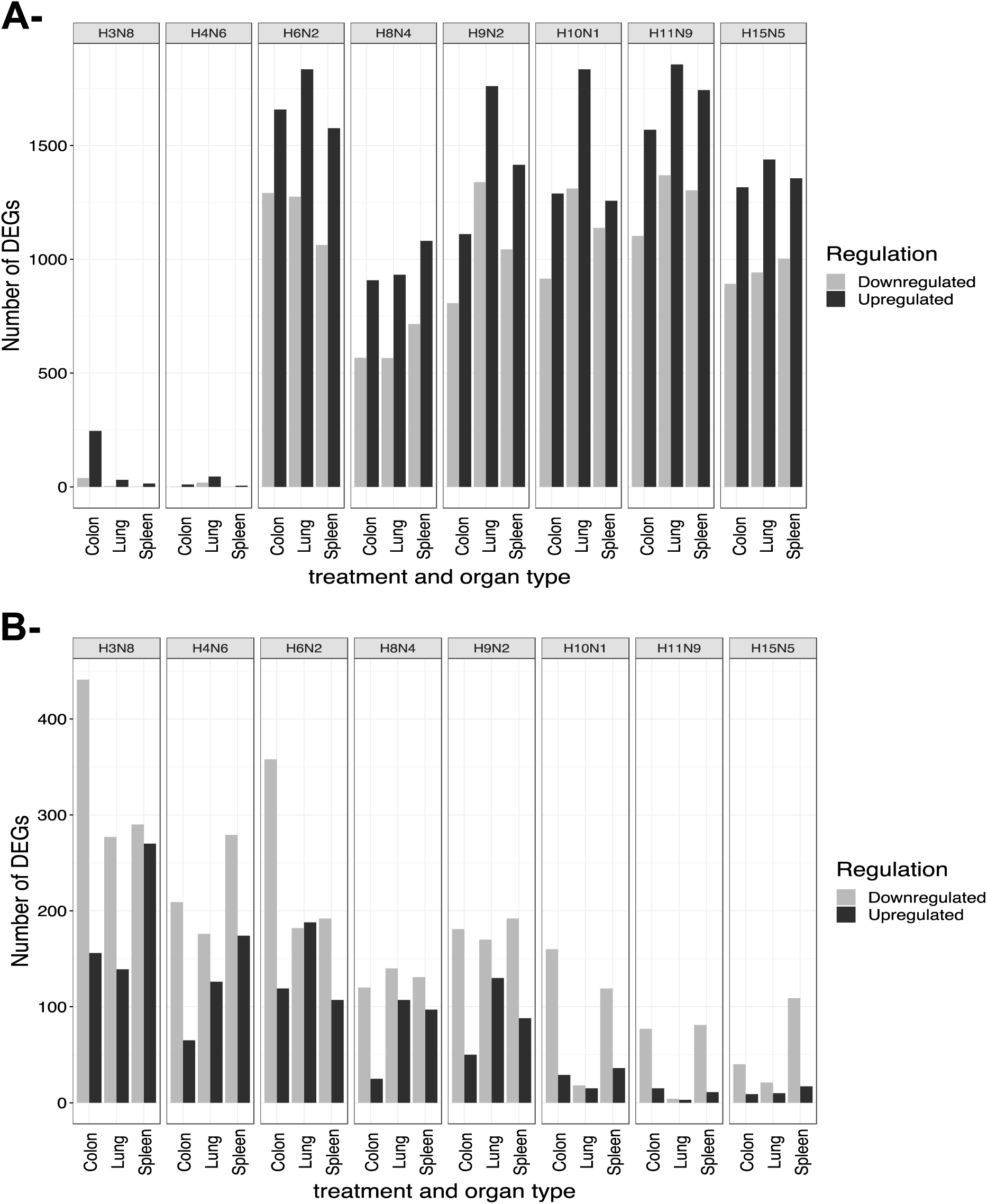
Differentially expressed genes in infected chickens (a) and tufted ducks (b). Number of differentially expressed genes (DEGs) in each tissue type for each virus subtype in chicken (a) and tufted ducks (b). Genes with an adjusted p-value (FDR) of ≤ 0.05 that were up- or down regulated at least 10% were considered differentially expressed.

**Figure S5.**
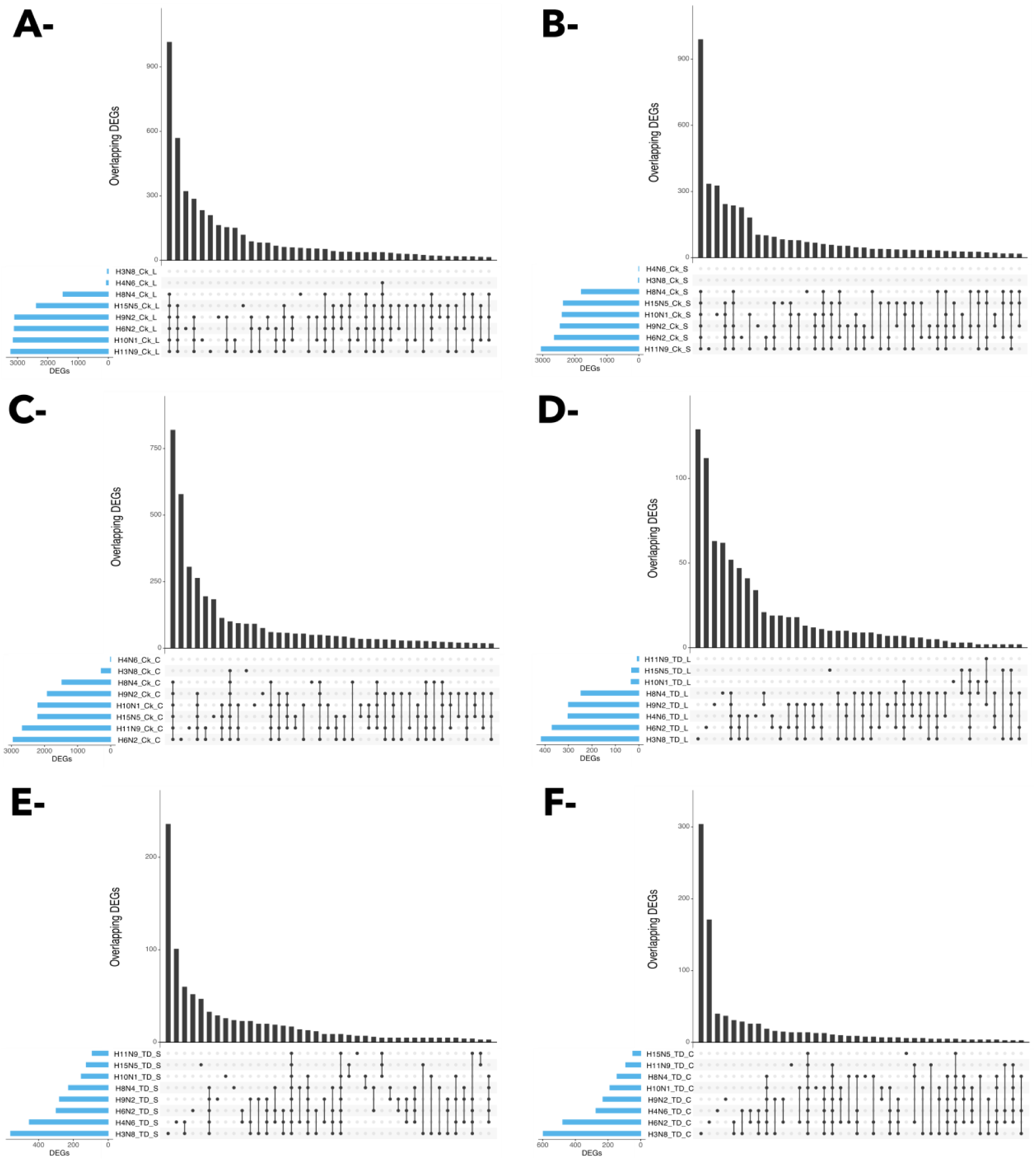
UpSet plot of the intersection of differentially expressed genes in chickens (a-c) and tufted ducks (d-e) tissues displayed for each virus subtype. Ck = chickens, TD= tufted ducks, L = lung, S = spleen C = colon. The nature of a given intersection is indicated by the dots below the bar plot. For instance, the genes in the 9^th^ column in A are differentially expressed in chicken lung following treatment with H10N1 and H11N9, whereas the genes in the 3^rd^ column in A were differentially expressed in chicken lung in response to H6N2 only.

**Figure S6.**
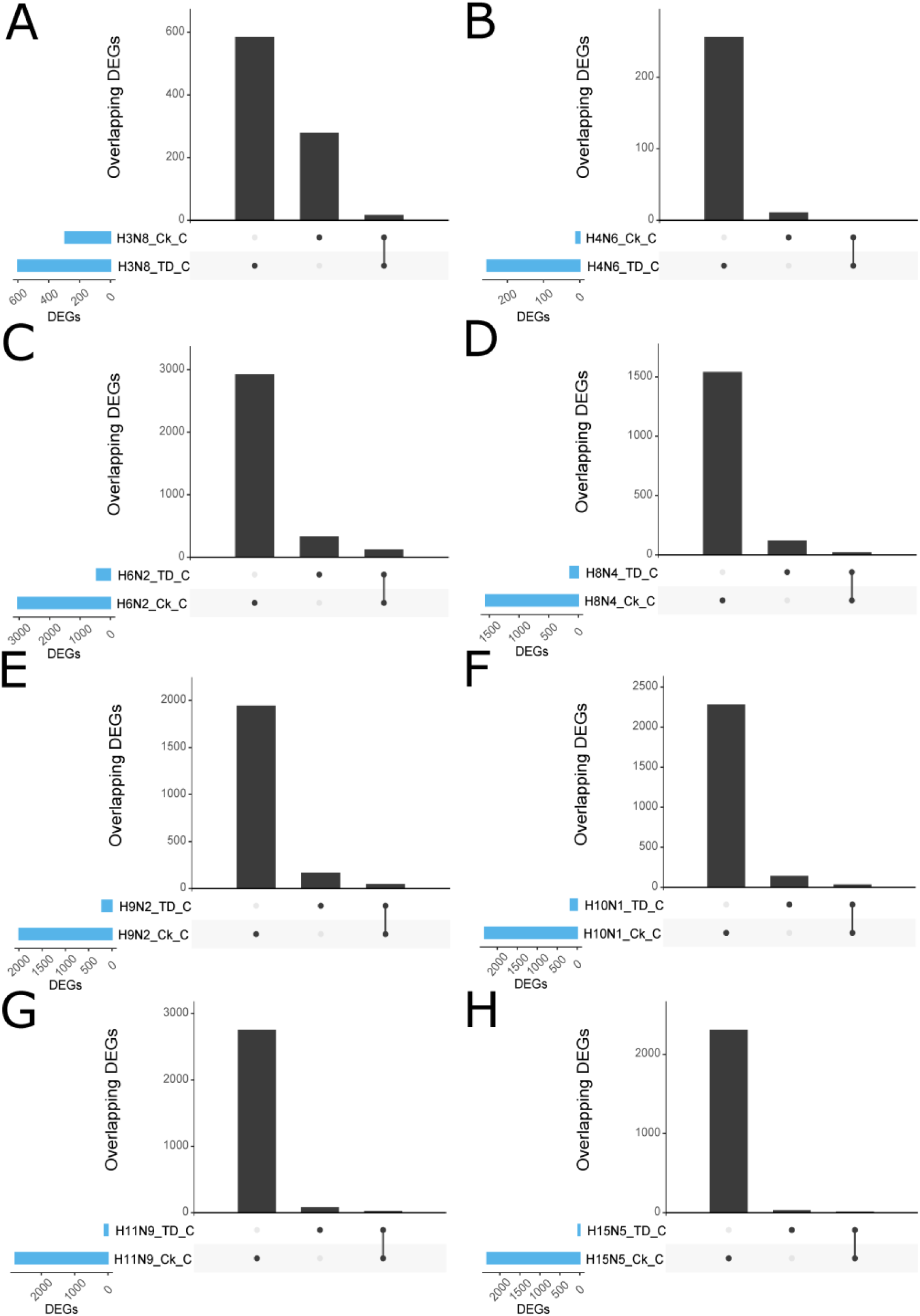
UpSet plot of the intersection of differentially expressed orthologous genes in tufted duck and chicken colon tissue, dispalyed for each virus subtype. A) H3N8, B) H4N6, C) H6N2, D) H8N4, E) H9N2, F) H10N1, G) H11N9, and H) H15N5. Ck = chickens, TD= tufted ducks. The nature of a given intersection is indicated by the dots below the bar plot. For instance, the genes in the first column in A are differentially expressed in tufted duck (TD) but not in chicken (Ck). Orthologs represent reciprocal best blast hits between the two species.

**Figure S7.**
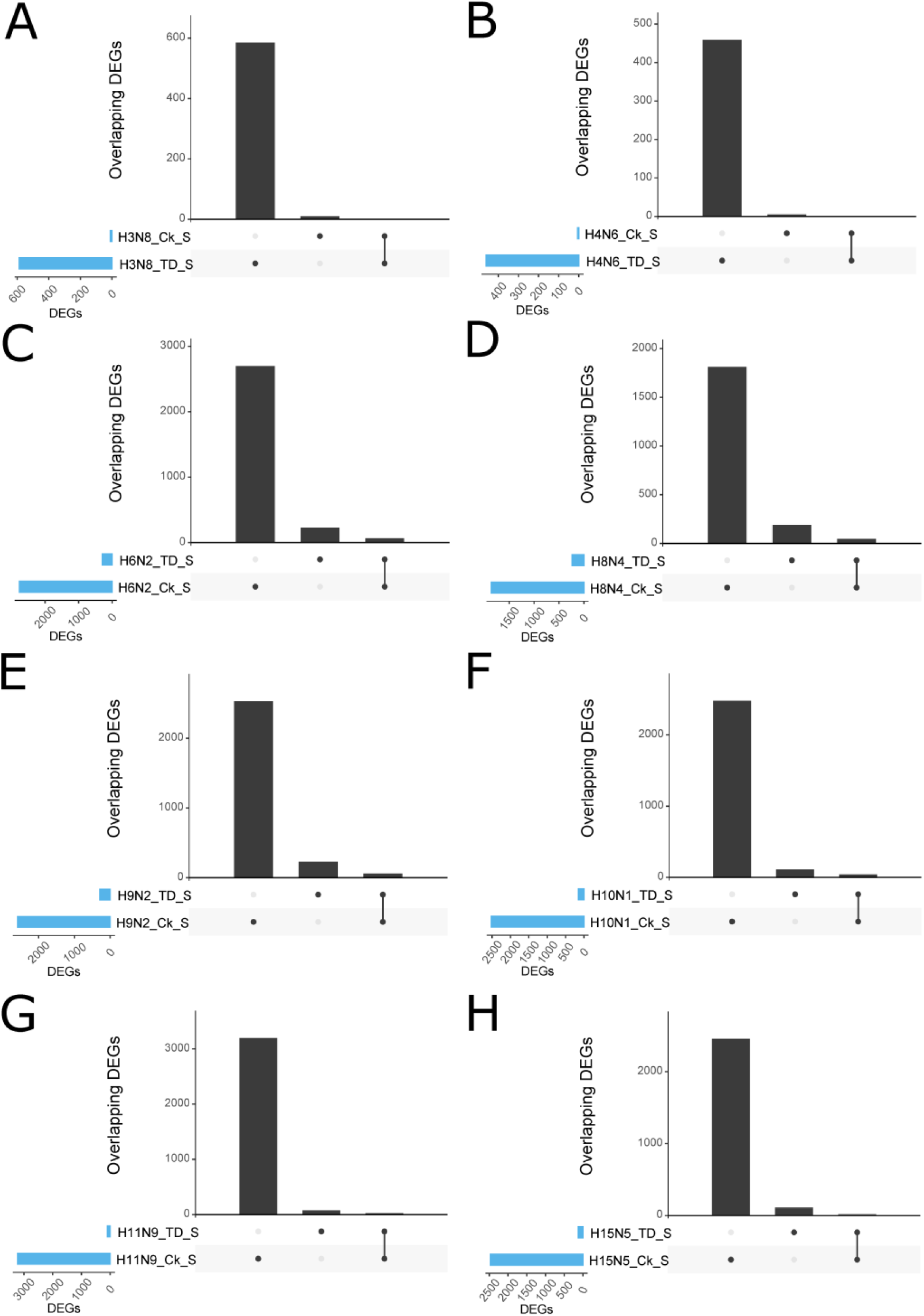
UpSet plot of the intersection of differentially expressed orthologous genes in tufted duck and chicken spleen tissue, displayed for each virus subtype. A) H3N8, B) H4N6, C) H6N2, D) H8N4, E) H9N2, F) H10N1, G) H11N9, and H) H15N5. Ck = chickens, TD= tufted ducks. The nature of a given intersection is indicated by the dots below the bar plot. For instance, the genes in the first column in A are differentially expressed in tufted duck (TD) but not in chicken (Ck). Orthologs represent reciprocal best blast hits between the two species.

**Figure S8.**
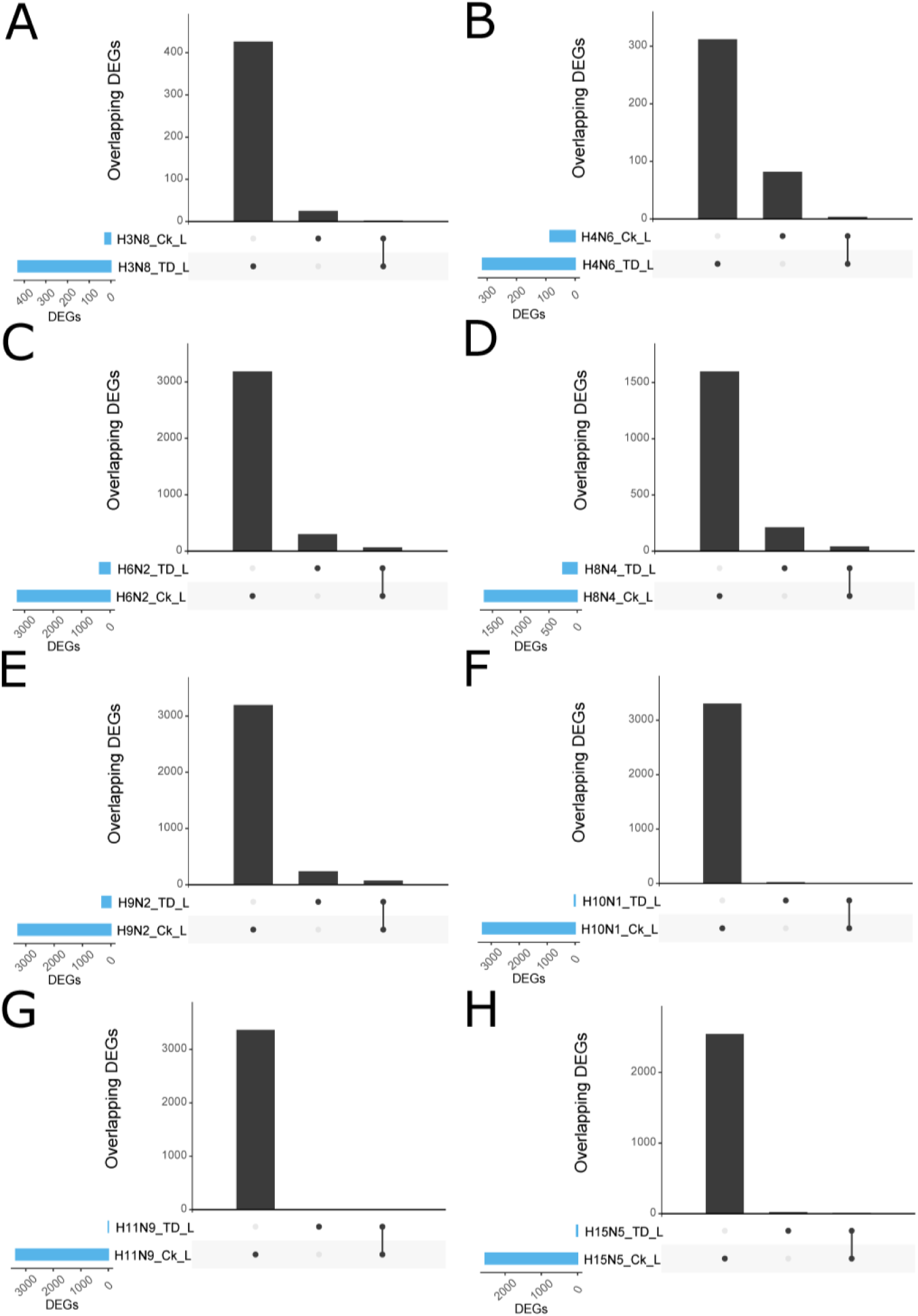
UpSet plot of the intersection of differentially expressed orthologous genes in tufted duck and chicken lung tissue, displayed for each virus subtype. A) H3N8, B) H4N6, C) H6N2, D) H8N4, E) H9N2, F) H10N1, G) H11N9, and H) H15N5. Ck = chickens, TD= tufted ducks. The nature of a given intersection is indicated by the dots below the bar plot. For instance, the genes in the first column in A are differentially expressed in tufted duck (TD) but not in chicken (Ck). Orthologs represent reciprocal best blast hits between the two species.

**Figure S9.**
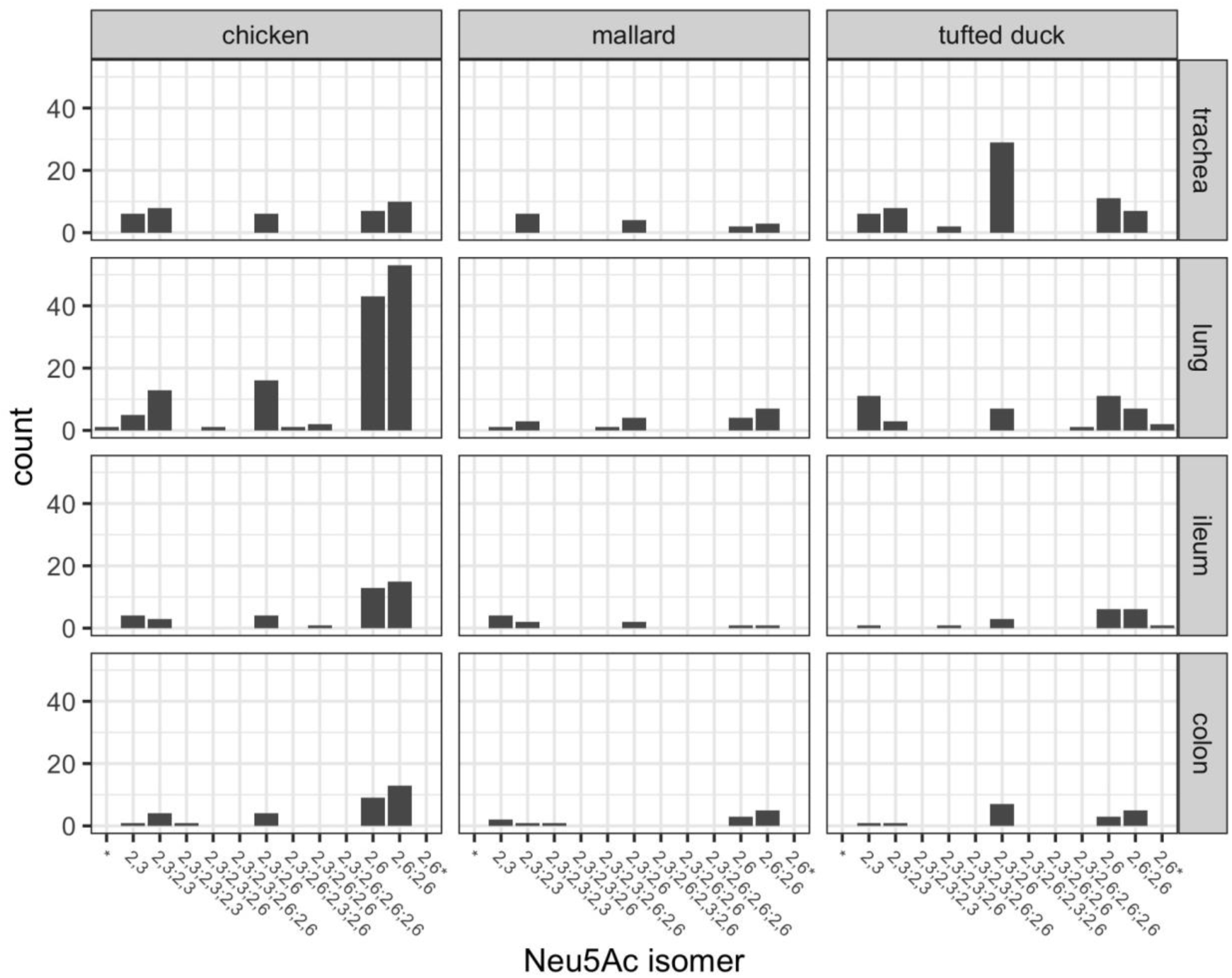
Distribution of sialylated glycoforms in chicken, mallard, and tufted duck. The figure illustrates the frequency of the different sialylated glycoforms (mono- to polysialylated glycoforms) identified in chicken, mallard, and tufted duck trachea, lung, ileum, and colon. 2,3 indicates α2,3-linked Neu5Ac and 2,6 indicates α2,6-linked Neu5Ac. A separation with; indicates that the glycan carries multiple Neu5Acs with different linkages. * indicates undetermined.

**Figure S10.**
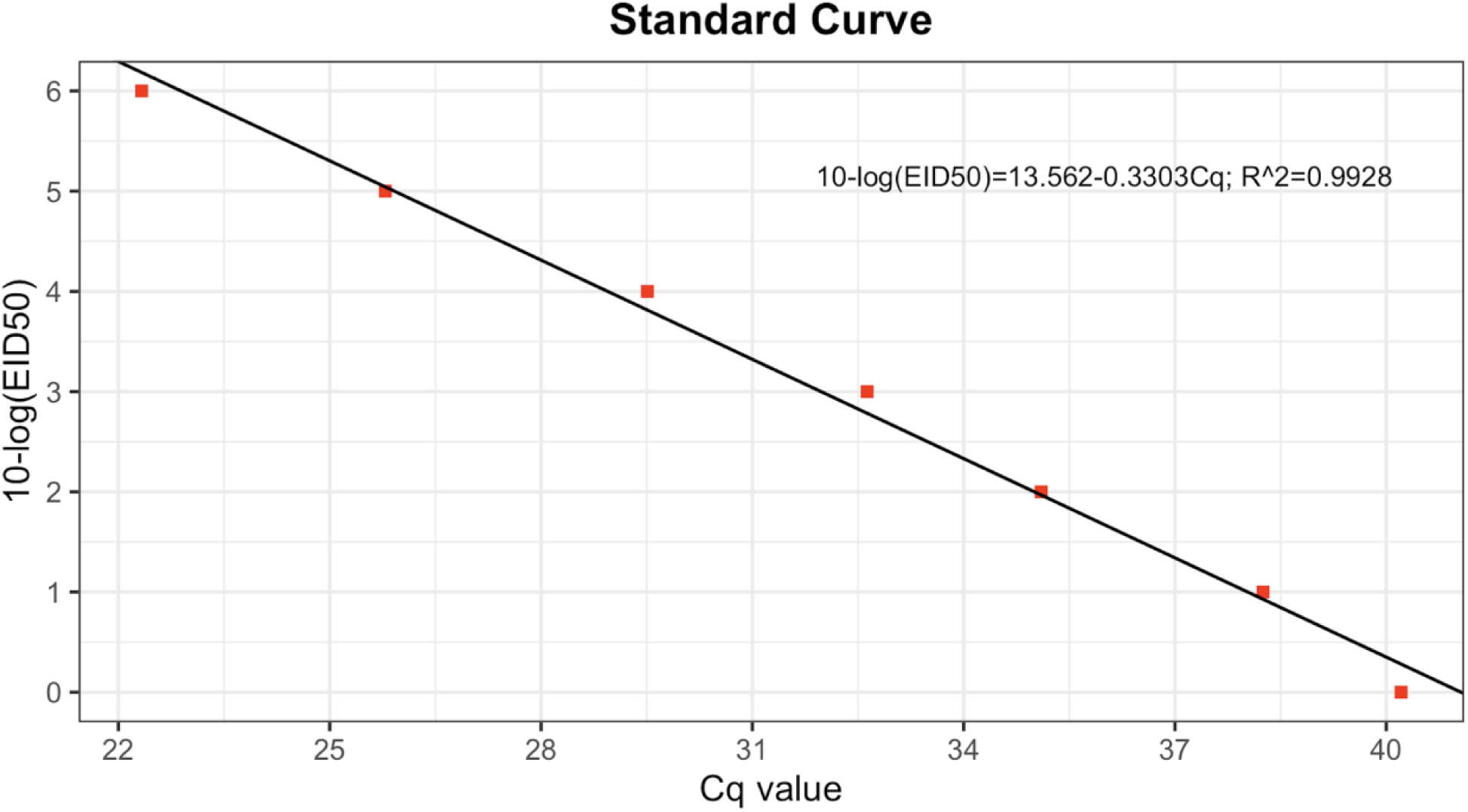
Rt-qPCR standard curve for the titeration of the inoculum. Standard curve was obtained from three 10-fold dilutions of the inoculum. Red squares showing the Cq values. Correlation coefficient *R*^2^ (0.998)

### Supplementary tables

**Table S1.** Investigated virus isolates.

| Subtype | Virus Designation |
| --- | --- |
| H3N8 | A/Mallard/Sweden/101487/2009 |
| H4N6 | A/Mallard/Sweden/80148/2008 |
| H6N2 | A/Mallard/Sweden/99825/2009 |
| H8N4 | A/Mallard/Sweden/58256/2006 |
| H9N2 | A/Mallard/Sweden/67860/2007 |
| H10N1 | A/Mallard/Sweden/102087/2009 |
| H11N9 | A/Mallard/Sweden/102103/2009 |
| H15N5 | A/Mallard/Sweden/139647/2012 |

**Table S2.**
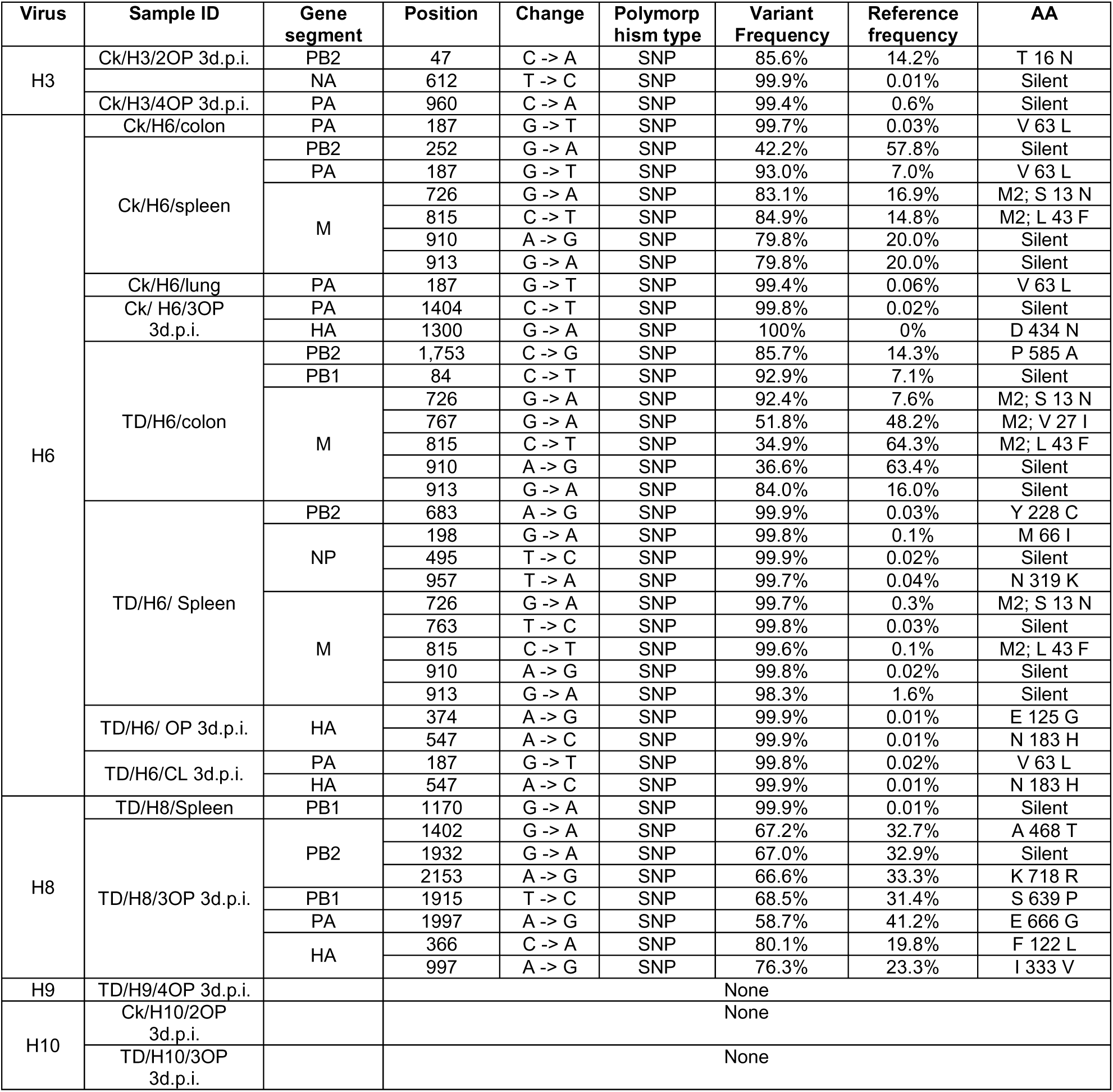
Variant detection after influenza A virus infection using virus inoculum as a reference.

**Table S3.**
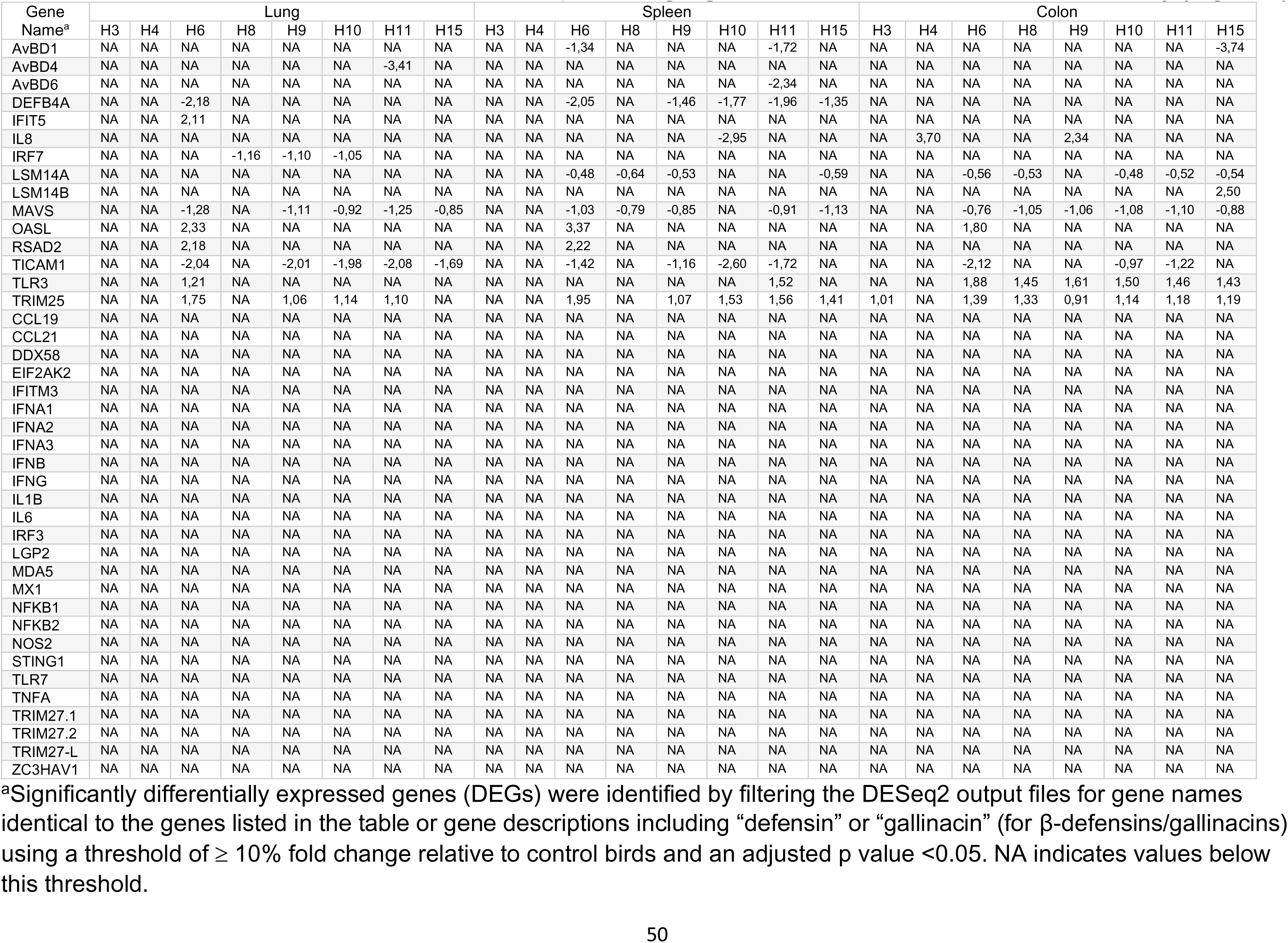
Genes used for construction of heat map showing significant DEGs related to innate immunity (Figure 7)

**Table S4.**
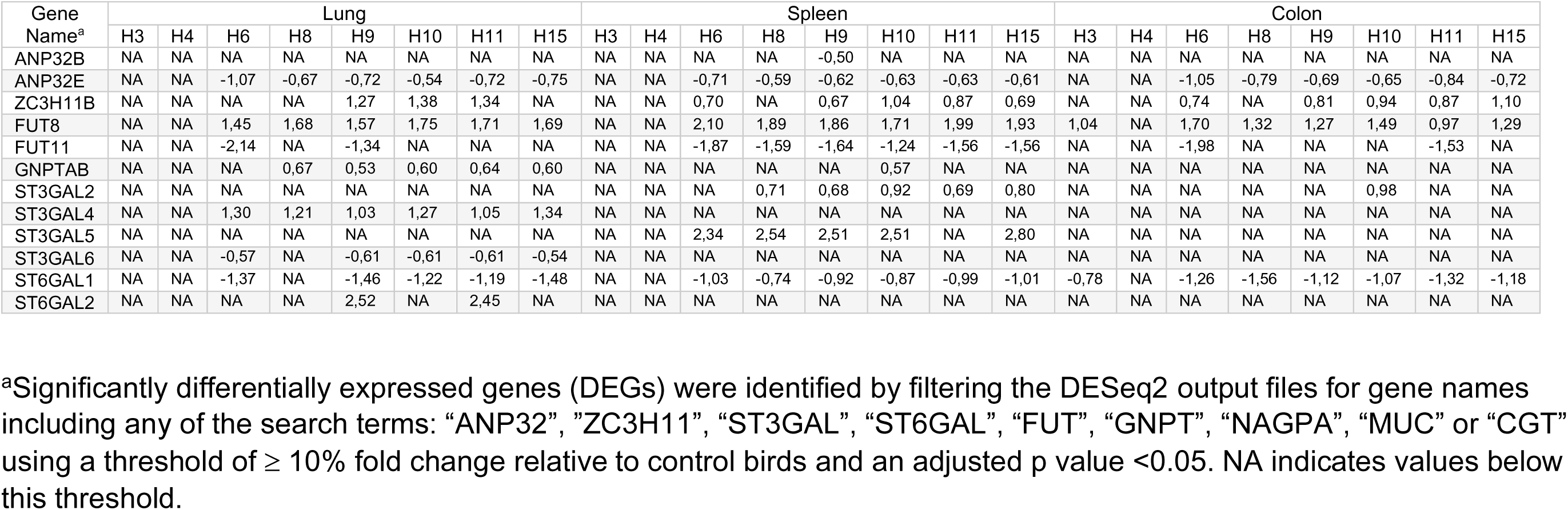
Differentially expressed genes identified by filtering DESeq2 output files from infected chickens using search terms^a^ related to glycosylation and transcription (Figure 8a)

**Table S5.** Differentially expressed genes identified by filtering DESeq2 output files from infected tufted ducks using search terms^a^ related to glycosylation and transcription (Figure 4b)

| Gene Name<sup>a</sup> | Lung |  |  |  |  |  |  |  | Spleen |  |  |  |  |  |  |  | Colon |  |  |  |  |  |  |  |
| --- | --- | --- | --- | --- | --- | --- | --- | --- | --- | --- | --- | --- | --- | --- | --- | --- | --- | --- | --- | --- | --- | --- | --- | --- |
|  | H3 | H4 | H6 | H8 | H9 | H10 | H11 | H15 | H3 | H4 | H6 | H8 | H9 | H10 | H11 | H15 | H3 | H4 | H6 | H8 | H9 | H10 | H11 | H15 |
| ZC3H11A | NA | NA | NA | NA | NA | NA | NA | NA | NA | NA | NA | NA | NA | NA | NA | NA | NA | NA | -1,42 | -1,10 | -1,13 | -1,12 | NA | NA |
| POFUT1 | NA | NA | NA | -2,42 | NA | NA | NA | NA | NA | NA | NA | NA | NA | NA | NA | NA | NA | NA | NA | NA | NA | NA | NA | NA |
| FUT4 | NA | NA | NA | NA | NA | NA | NA | NA | NA | -1,42 | NA | NA | NA | NA | NA | NA | NA | NA | NA | NA | NA | NA | NA | NA |
| FUT8 | NA | NA | NA | NA | NA | NA | NA | NA | NA | 0,92 | NA | 0,68 | NA | NA | NA | NA | NA | NA | NA | NA | NA | NA | NA | NA |
| FUT10 | NA | NA | 1,72 | NA | 1,62 | NA | NA | NA | NA | NA | NA | NA | NA | NA | NA | NA | NA | NA | NA | NA | NA | NA | NA | NA |
| ST3GAL2 | NA | NA | NA | NA | NA | NA | NA | NA | NA | NA | NA | NA | NA | NA | NA | NA | -0,91 | NA | NA | NA | NA | NA | NA | NA |
| ST6GALNAC2 | NA | NA | NA | NA | NA | NA | NA | NA | NA | NA | NA | NA | NA | NA | NA | NA | -1,73 | NA | NA | NA | NA | NA | NA | NA |
| ST6GALNAC4 | NA | NA | NA | NA | NA | NA | NA | NA | NA | NA | NA | NA | NA | NA | NA | NA | -1,91 | NA | NA | NA | NA | NA | NA | NA |
| GNPTAB | NA | NA | NA | NA | NA | NA | NA | NA | 0,94 | NA | NA | NA | 0,96 | NA | NA | NA | NA | NA | NA | NA | NA | NA | NA | NA |
| MUC2 | NA | NA | NA | NA | 3,96 | NA | NA | NA | 6,37 | NA | NA | NA | NA | 4,42 | NA | NA | NA | NA | NA | NA | NA | NA | NA | NA |

<sup>a</sup>Significantly differentially expressed genes (DEGs) were identified by filtering the DESeq2 output files for gene names including any of the search terms: “ANP32”, “ZC3H11”, “ST3GAL”, “ST6GAL”, “FUT”, “GNPT”, “NAGPA”, “MUC” or “CGT” using a threshold of $\geq 10\%$ fold change relative to control birds and an adjusted p value $<0.05$ . NA indicates values below this threshold.

**Table S6.**
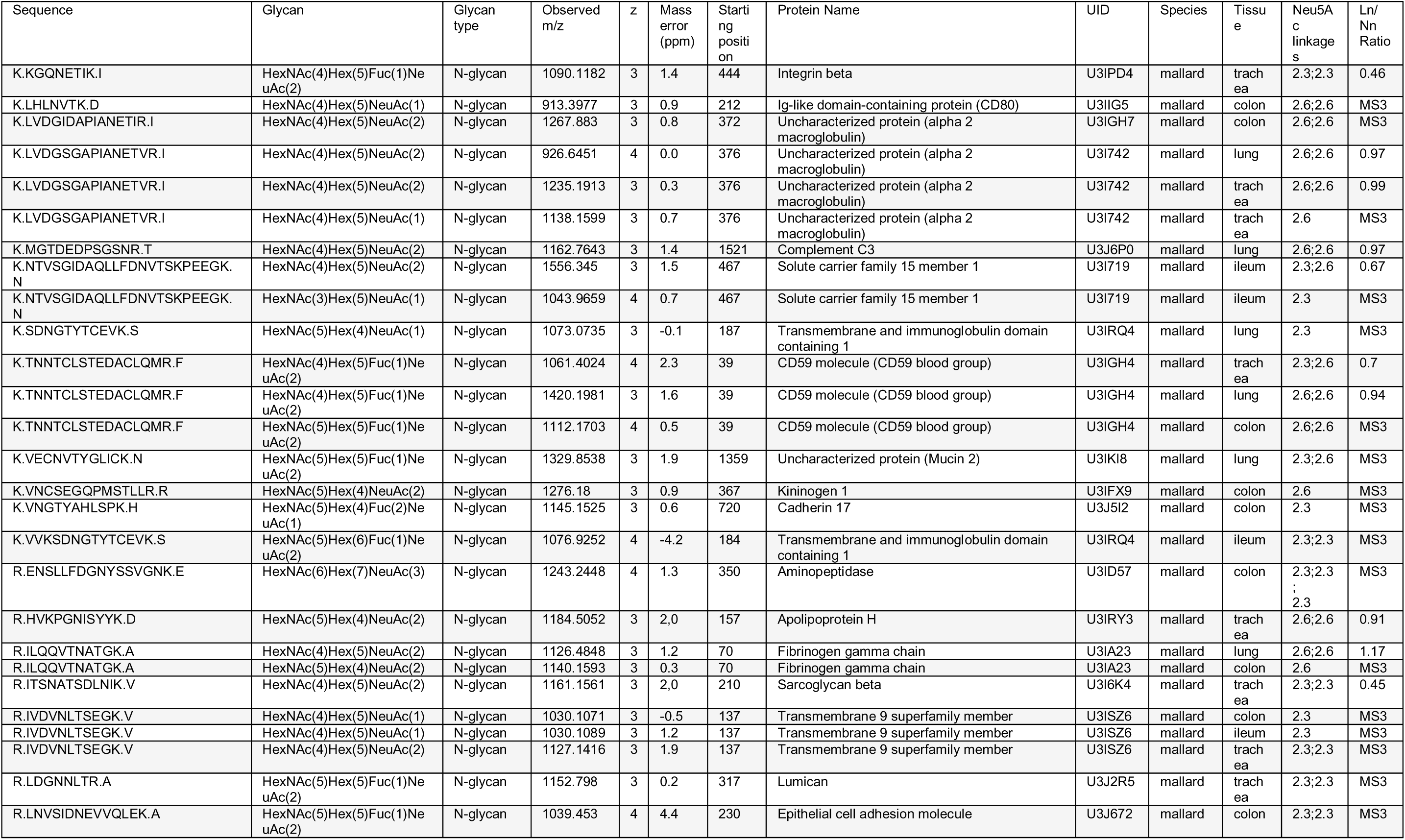
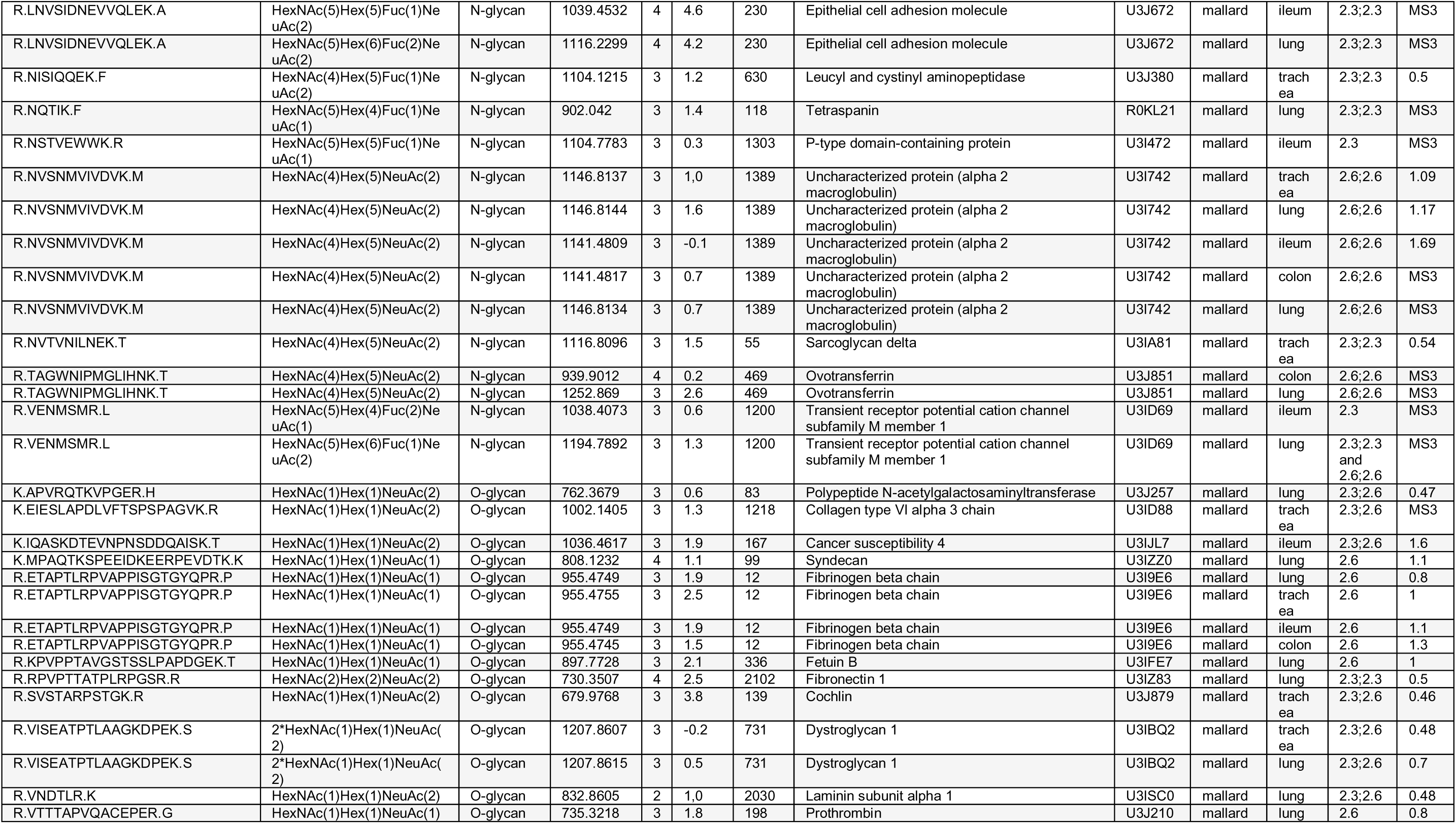
Annotated list of sialylated *N*- and *O*-linked glycans detected in mallard tissues. *m/z* — mass to charge ratio, z — charge, UID — Uniprot ID, and Ln/Nn — number of LacNAc/number of Neu5Ac ratio, 0.4–0.6 for Neu5Acα2,3 and 0.8–1.5 for Neu5Acα2,6 terminated glycopeptides.

**Table S7.**
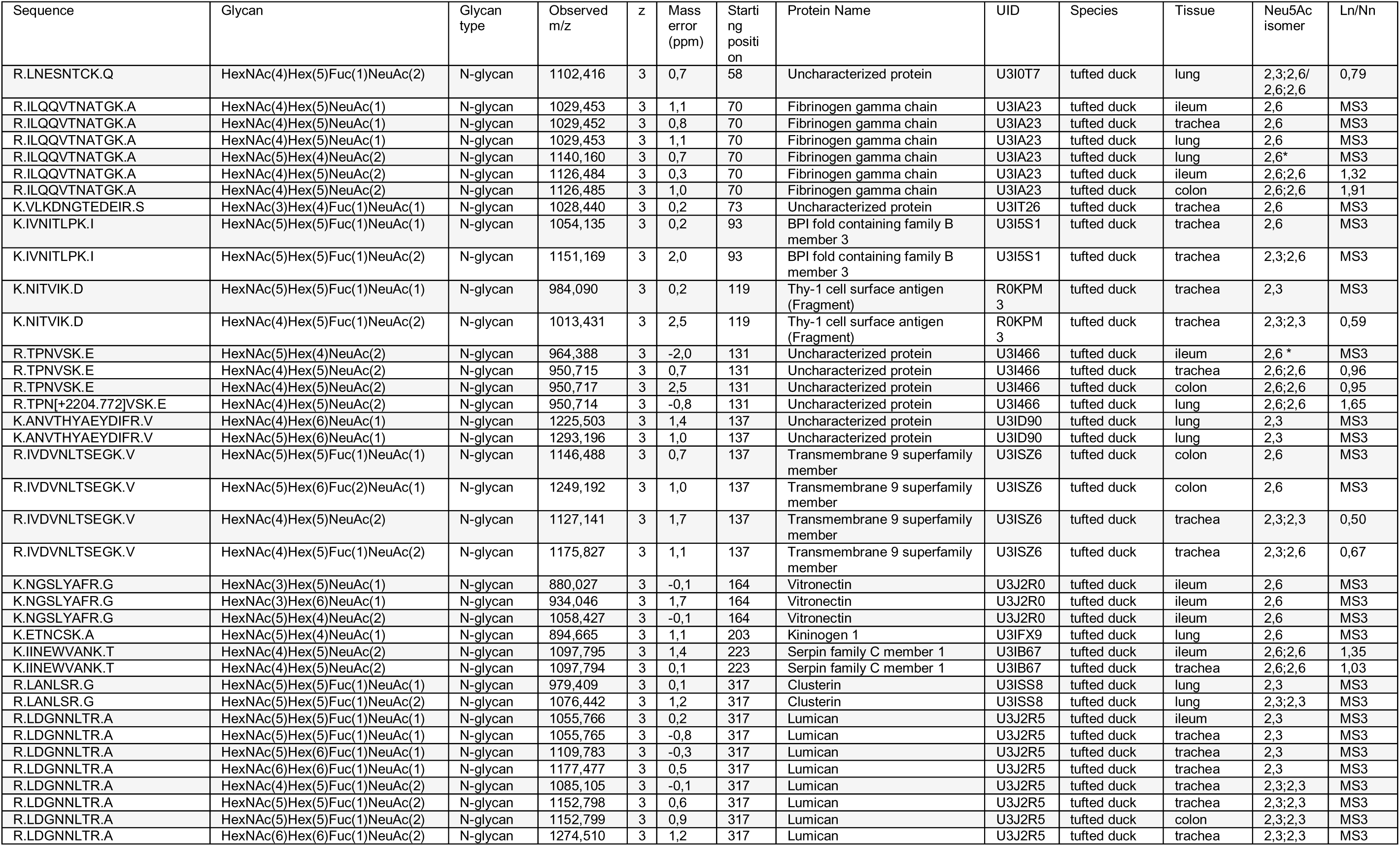
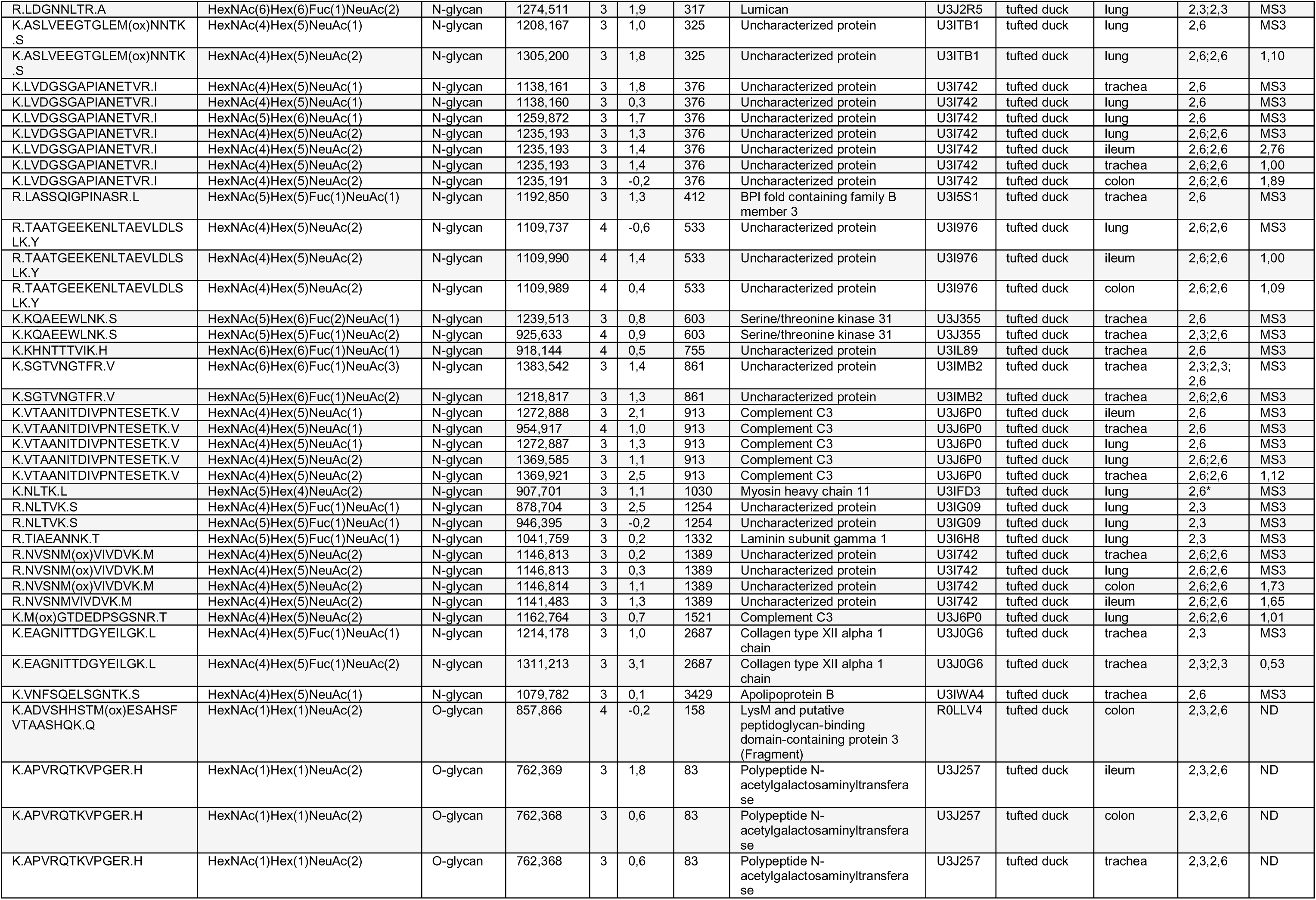
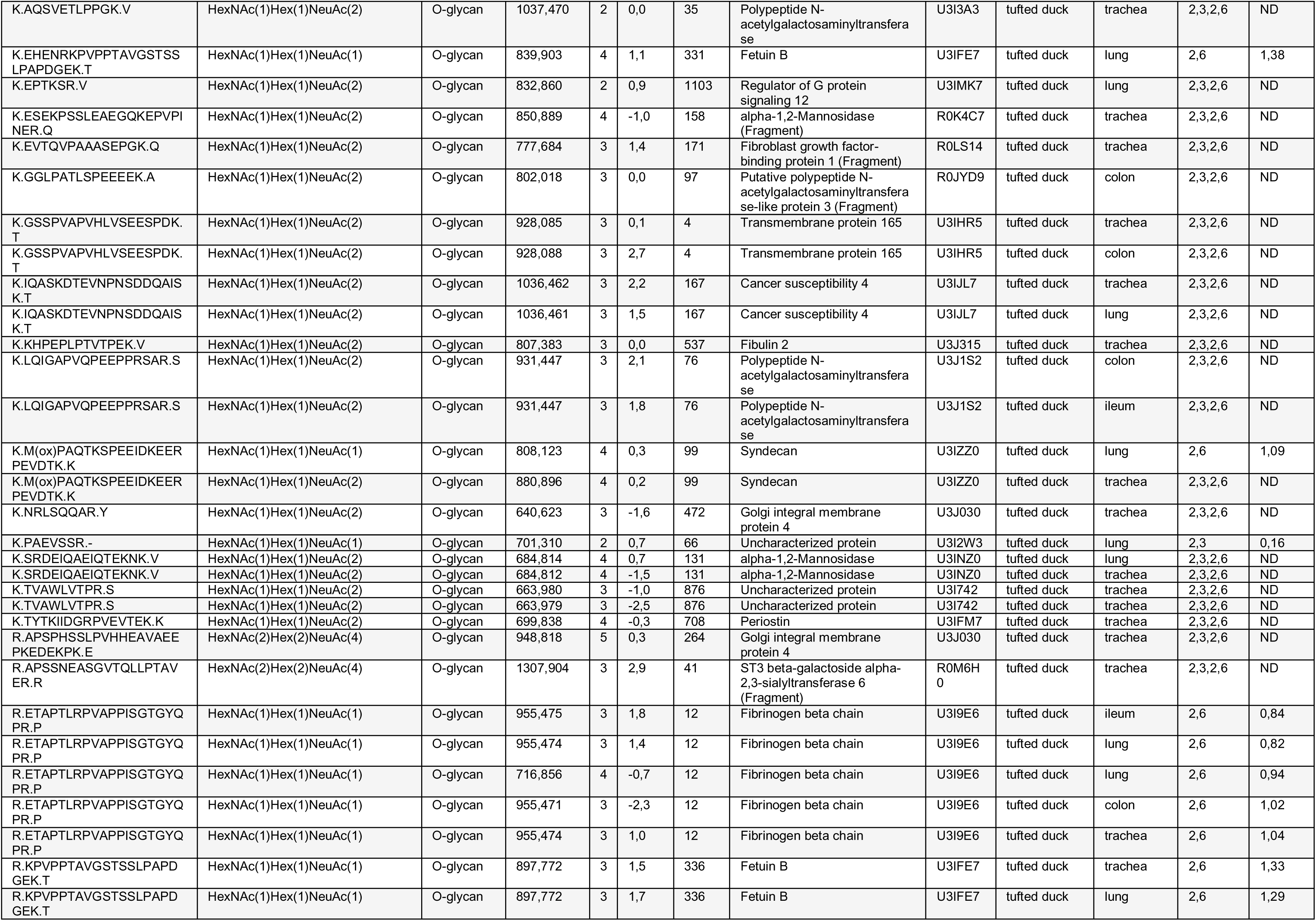
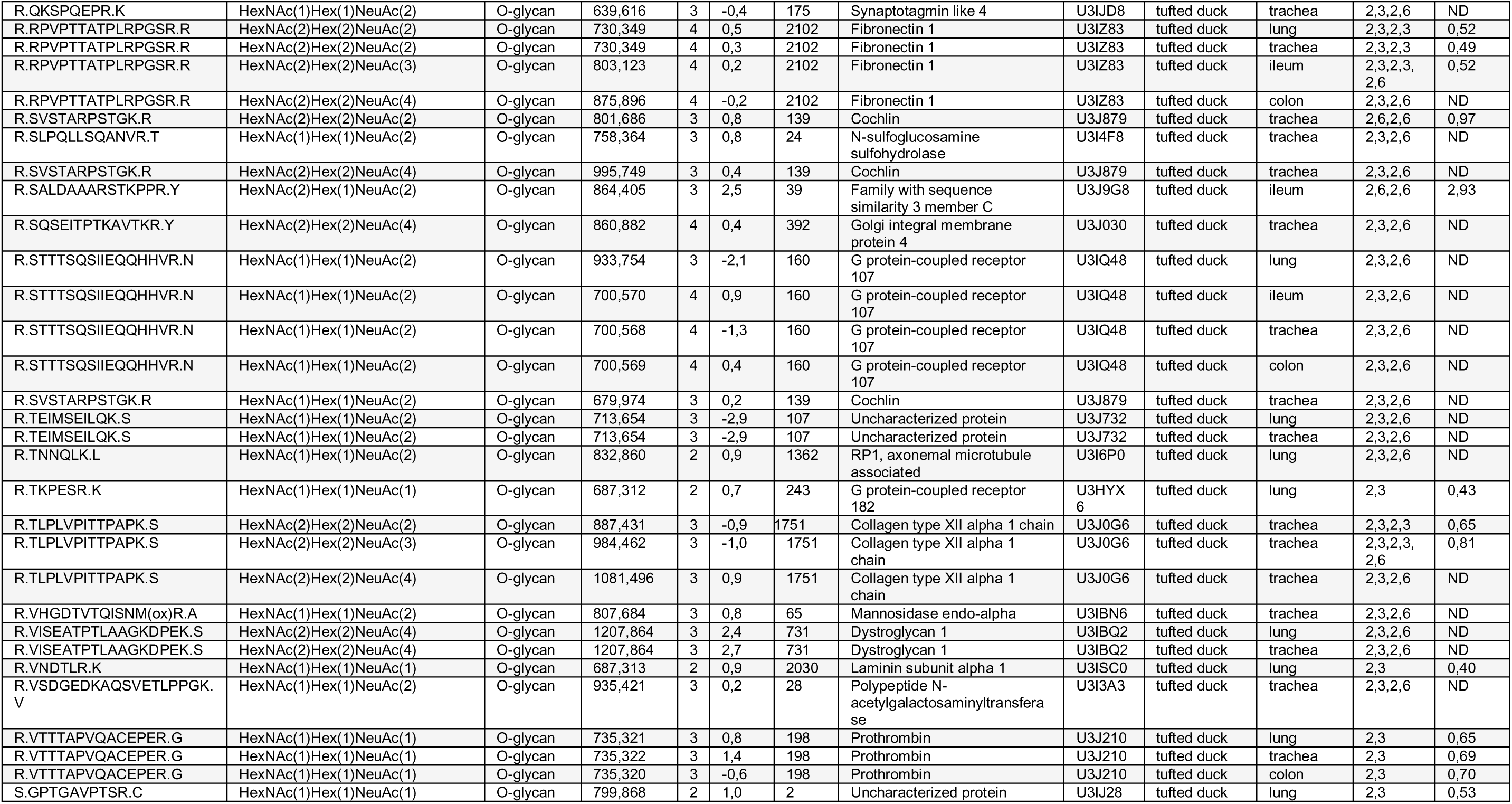
Annotated list of sialylated *N*- and *O*-linked glycans detected in sampled tufted duck tissues. *m/z* — mass to charge ratio, z — charge, UID — Uniprot ID, and Ln/Nn — number of LacNAc/number of Neu5Ac ratio, 0.4–0.6 for Neu5Acα2,3 and 0.8–1.5 for Neu5Acα2,6 terminated glycopeptides. ND= not determined. Disialo core 1 glycopeptides were assumed to be 2,3;2,6. * One NeuAc is on Gal and the second is on GalNAc.

**Table S8.**
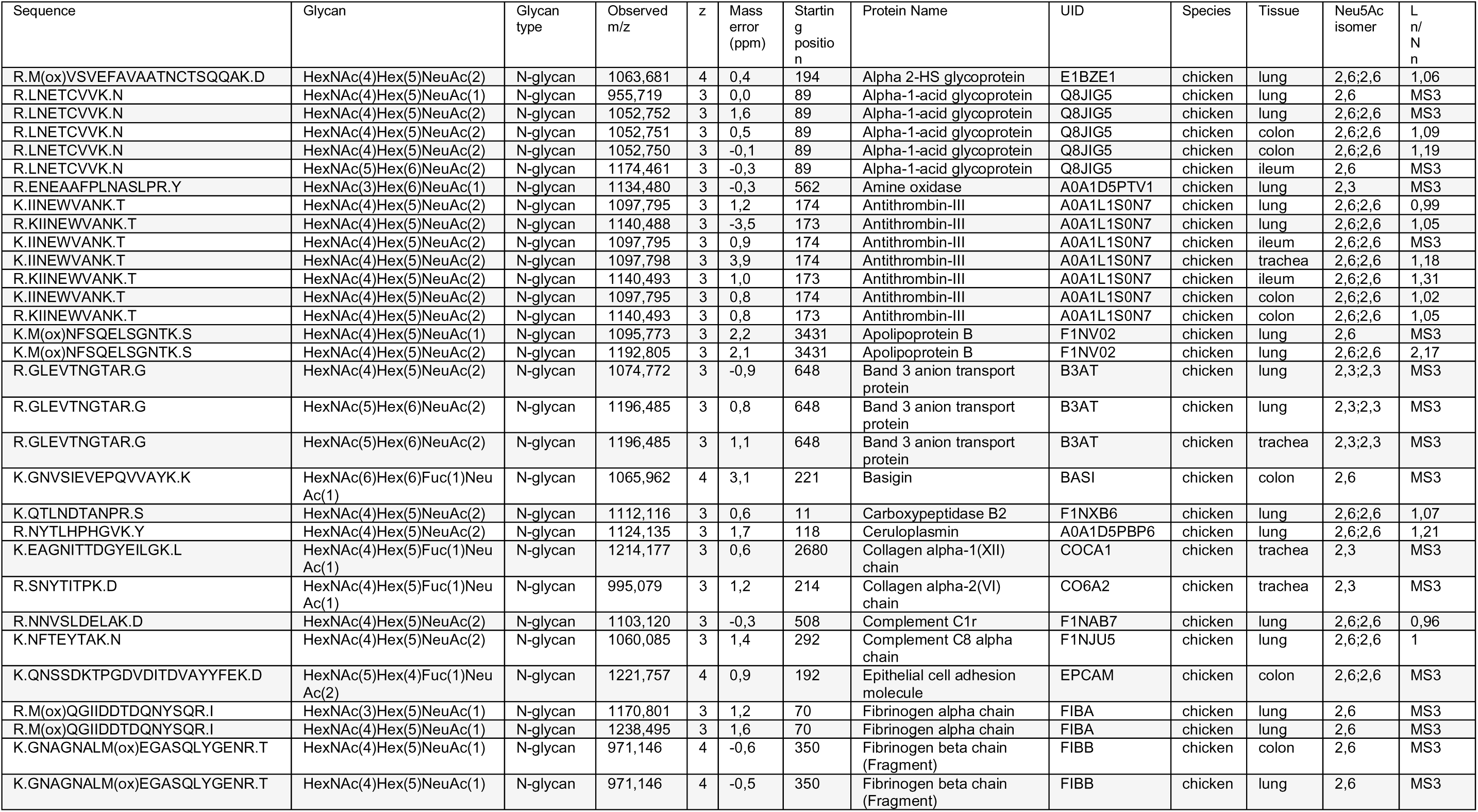
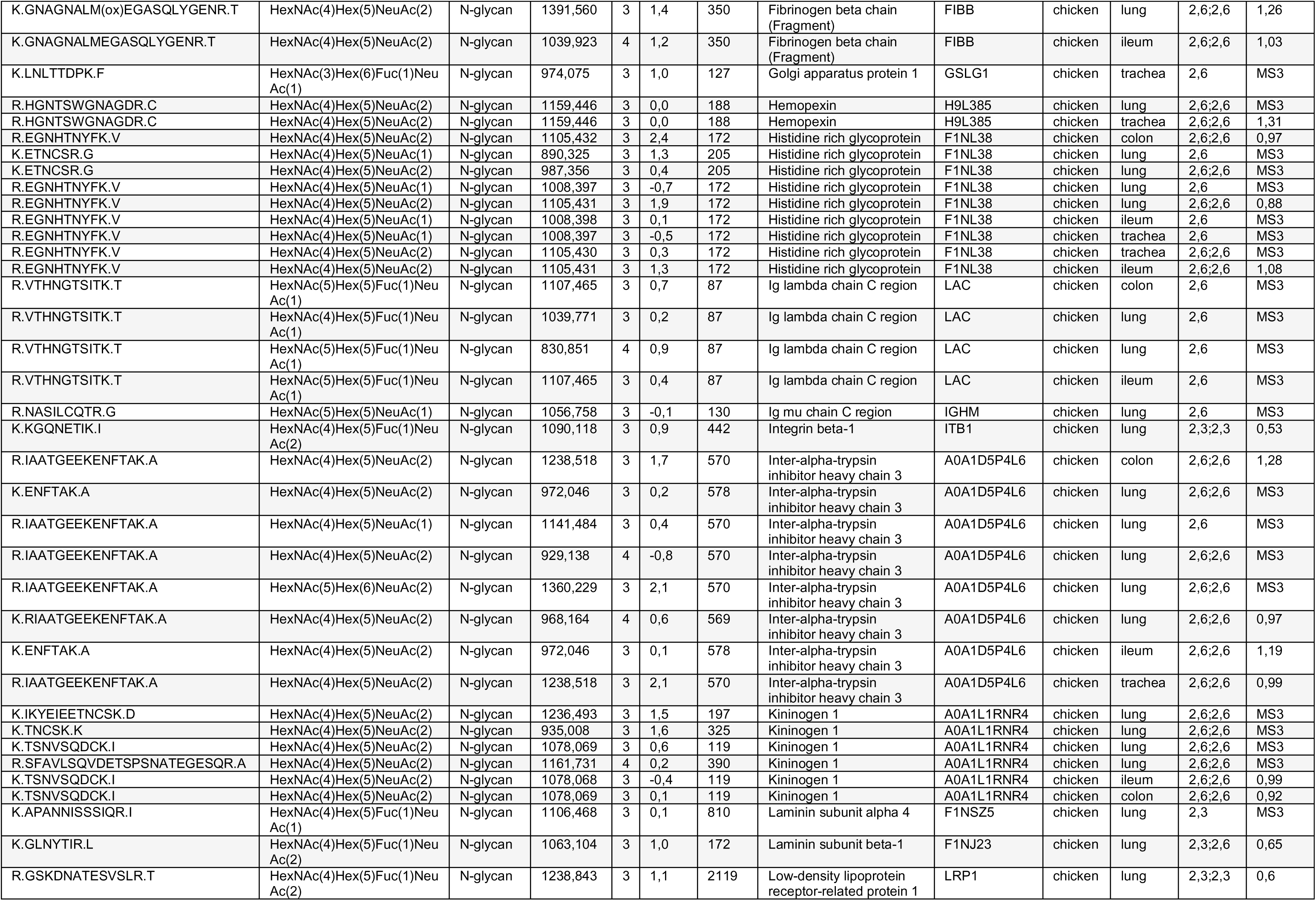
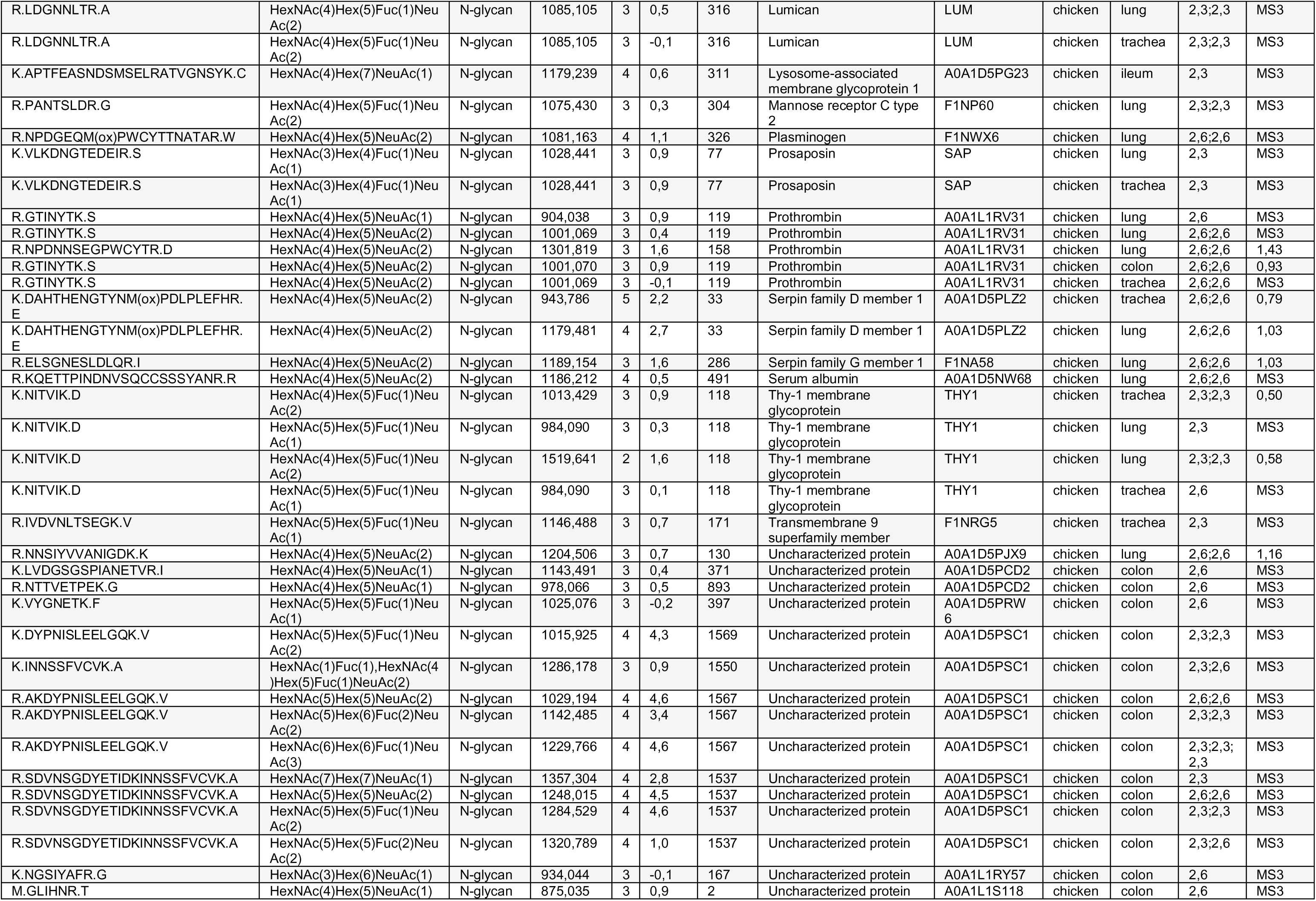
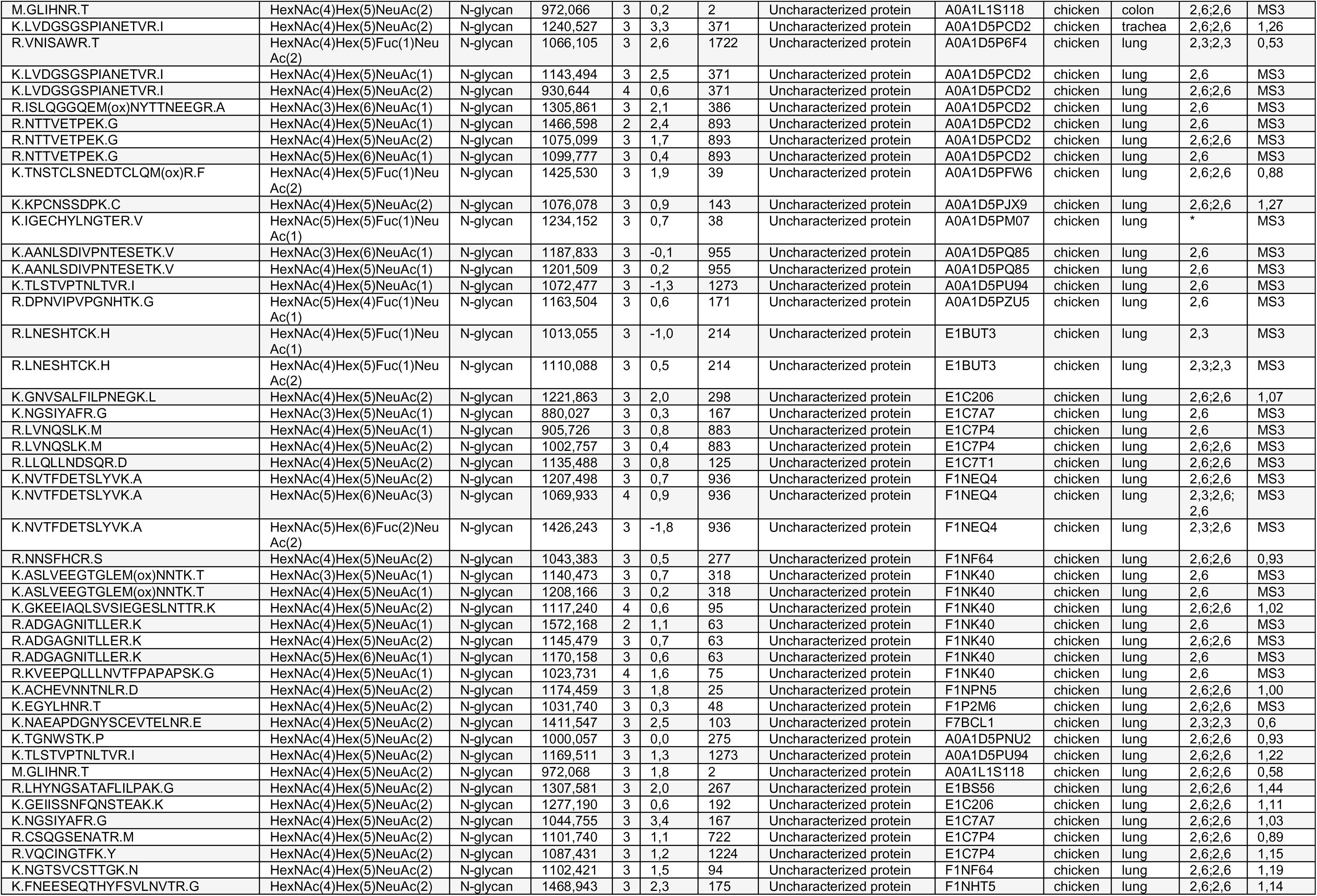
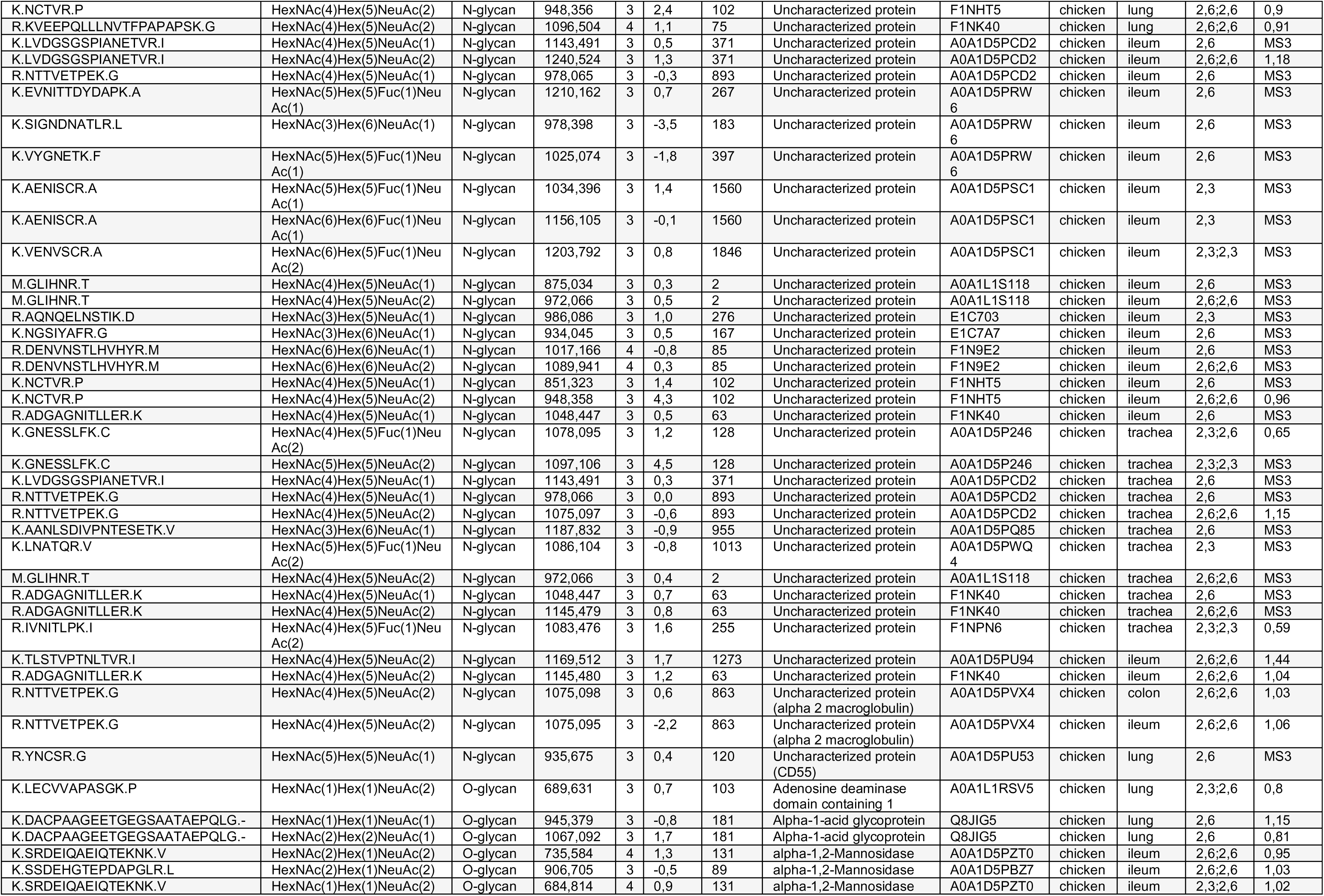
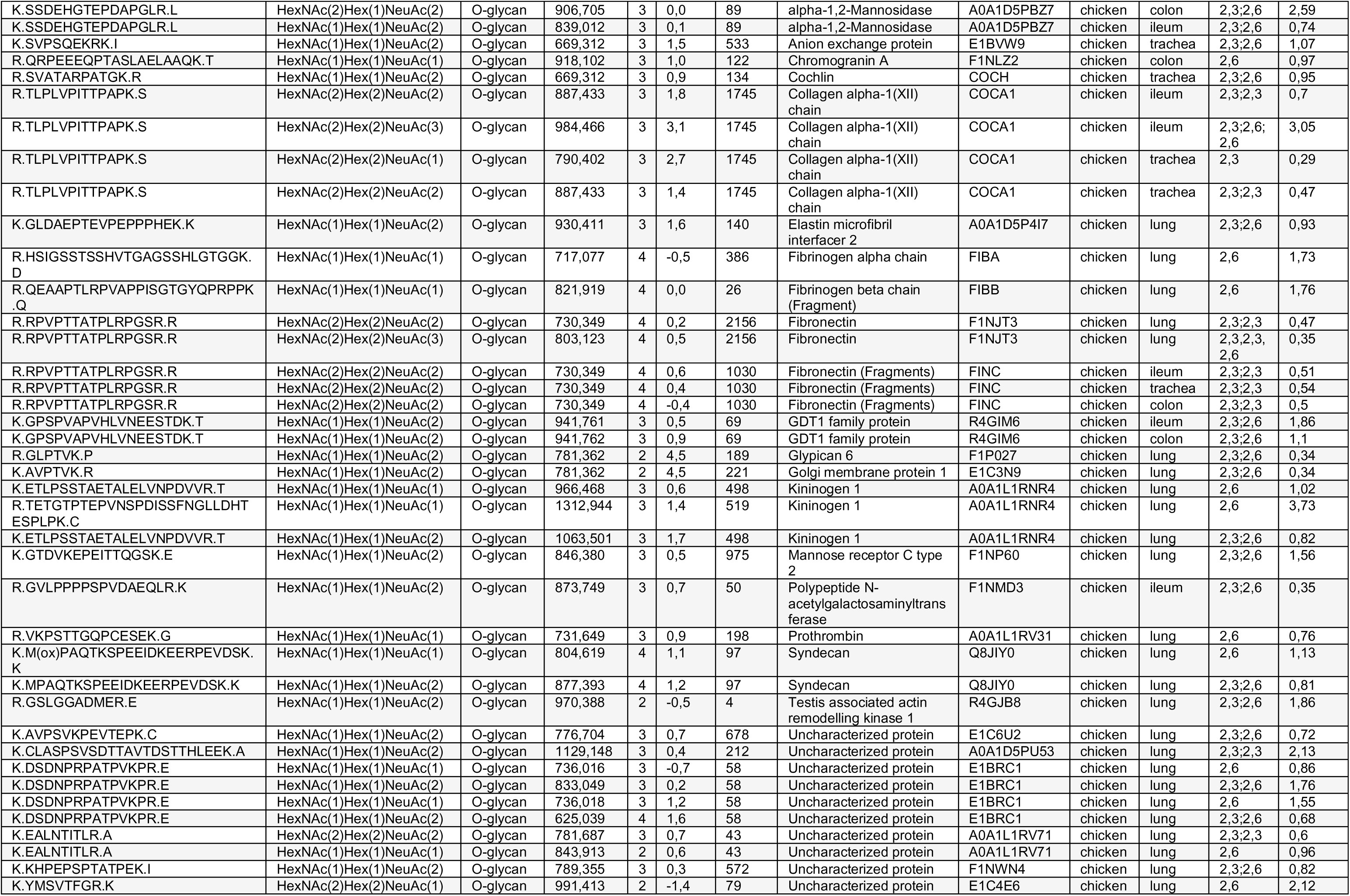
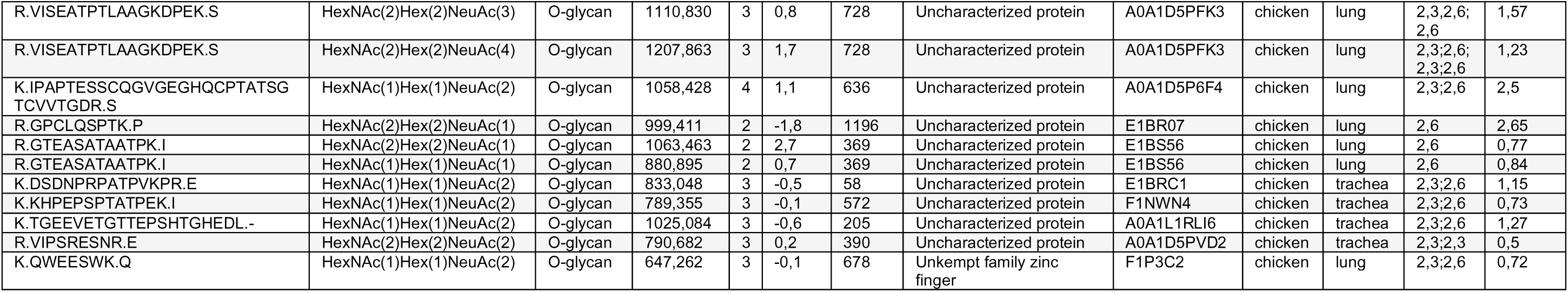
Annotated list of sialylated *N*- and *O*-linked glycans detected in sampled chicken tissues. *m/z* — mass to charge ratio, z — charge, UID — Uniprot ID, and Ln/Nn — number of LacNAc/number of Neu5Ac ratio, 0.4–0.6 for Neu5Acα2,3 and 0.8–1.5 for Neu5Acα2,6 terminated glycopeptides.

## References

1. Kreuder Johnson, C. et al. Spillover and pandemic properties of zoonotic viruses with high host plasticity. Sci Rep 5, 14830–14830, doi:10.1038/srep14830 (2015).

2. Yen, H. L. & Webster, R. G. Pandemic influenza as a current threat. Curr Top Microbiol Immunol 333, 3–24, doi:10.1007/978-3-540-92165-3_1 (2009).

3. Katz, J. M. et al. The public health impact of avian influenza viruses. Poult Sci 88, 872–879, doi:10.3382/ps.2008-00465 (2009).

4. Horimoto, T. & Kawaoka, Y. Pandemic threat posed by avian influenza A viruses. Clin Microbiol Rev 14, 129–149, doi:10.1128/CMR.14.1.129-149.2001 (2001).

5. Short, K. R. et al. One health, multiple challenges: The inter-species transmission of influenza A virus. One Health 1, 1–13, doi:10.1016/j.onehlt.2015.03.001 (2015).

6. Subbarao, K. The Critical Interspecies Transmission Barrier at the Animal⁻Human Interface. Trop Med Infect Dis 4, 72, doi:10.3390/tropicalmed4020072 (2019).

7. Gottschalk, A. Carbohydrate residue of a urine mucoprotein inhibiting influenza virus haemagglutination. Nature 170, 662–663 (1952).

8. Gottschalk, A. The influenza virus enzyme and its mucoprotein substrate. Yale J Biol Med 26, 352–364 (1954).

9. Blix, F. G., Gottschalk, A. & Klenk, E. Proposed nomenclature in the field of neuraminic and sialic acids. Nature 179, 1088 (1957).

10. Rogers, G. N. & Paulson, J. C. Receptor determinants of human and animal influenza virus isolates: differences in receptor specificity of the H3 hemagglutinin based on species of origin. Virology 127, 361–373 (1983).

11. Rogers, G. N. & D’Souza, B. L. Receptor binding properties of human and animal H1 influenza virus isolates. Virology 173, 317–322 (1989).

12. Gambaryan, A. et al. Receptor specificity of influenza viruses from birds and mammals: new data on involvement of the inner fragments of the carbohydrate chain. Virology 334, 276–283, doi:10.1016/j.virol.2005.02.003 (2005).

13. Gambaryan, A. S. et al. Receptor-binding properties of influenza viruses isolated from gulls. Virology 522, 37–45, doi:10.1016/j.virol.2018.07.004 (2018).

14. Stevens, J., Blixt, O., Paulson, J. C. & Wilson, I. A. Glycan microarray technologies: tools to survey host specificity of influenza viruses. Nat Rev Microbiol 4, 857–864, doi:10.1038/nrmicro1530 (2006).

15. Gambaryan, A. S. et al. 6-sulfo sialyl Lewis X is the common receptor determinant recognized by H5, H6, H7 and H9 influenza viruses of terrestrial poultry. Virology journal 5, 85, doi:10.1186/1743-422x-5-85 (2008).

16. Walther, T. et al. Glycomic analysis of human respiratory tract tissues and correlation with influenza virus infection. PLoS Pathog 9, e1003223, doi:10.1371/journal.ppat.1003223 (2013).

17. Shinya, K. et al. Avian flu: influenza virus receptors in the human airway. Nature 440, 435–436, doi:10.1038/440435a (2006).

18. Ellström, P., Jourdain, E., Gunnarsson, O., Waldenstrom, J. & Olsen, B. The “human influenza receptor” Neu5Ac alpha 2,6Gal is expressed among different taxa of wild birds. Arch Virol 154, 1533–1537, doi:10.1007/s00705-009-0476-8 (2009).

19. Liu, Y. et al. Influenza A virus receptors in the respiratory and intestinal tracts of pigeons. Avian Pathol 38, 263–266, doi:10.1080/03079450903055363 (2009).

20. Yu, J. E. et al. Expression patterns of influenza virus receptors in the respiratory tracts of four species of poultry. Journal of Veterinary Science 12, 7, doi:10.4142/jvs.2011.12.1.7 (2011).

21. Costa, T. et al. Distribution patterns of influenza virus receptors and viral attachment patterns in the respiratory and intestinal tracts of seven avian species. Vet Res 43, 28, doi:10.1186/1297-9716-43-28 (2012).

22. Franca, M., Stallknecht, D. E. & Howerth, E. W. Expression and distribution of sialic acid influenza virus receptors in wild birds. Avian Pathol 42, 60–71, doi:10.1080/03079457.2012.759176 (2013).

23. Eriksson, P. et al. Attachment Patterns of Human and Avian Influenza Viruses to Trachea and Colon of 26 Bird Species–Support for the Community Concept. Frontiers in Microbiology 10, 815 (2019).

24. Wille, M., Brojer, C., Lundkvist, A. & Jarhult, J. D. Alternate routes of influenza A virus infection in Mallard (Anas platyrhynchos). Vet Res 49, 110, doi:10.1186/s13567-018-0604-0 (2018).

25. Jourdain, E. et al. The pattern of influenza virus attachment varies among wild bird species. PLoS One 6, e24155, doi:10.1371/journal.pone.0024155 (2011).

26. Bergervoet, S. A. et al. Susceptibility of Chickens to Low Pathogenic Avian Influenza (LPAI) Viruses of Wild Bird- and Poultry-Associated Subtypes. Viruses 11, doi:10.3390/v11111010 (2019).

27. Bröjer, C. et al. Pathology of natural highly pathogenic avian influenza H5N1 infection in wild tufted ducks (Aythya fuligula). Journal of veterinary diagnostic investigation: official publication of the American Association of Veterinary Laboratory Diagnosticians, Inc 21, 579–587, doi:10.1177/104063870902100501 (2009).

28. Bröjer, C. et al. Pathobiology and virus shedding of low-pathogenic avian influenza virus (A/H1N1) infection in mallards exposed to oseltamivir. Journal of wildlife diseases 49, 103–113, doi:10.7589/2011-11-335 (2013).

29. Gillman, A. et al. Influenza A(H7N9) Virus Acquires Resistance-Related Neuraminidase I222T Substitution When Infected Mallards Are Exposed to Low Levels of Oseltamivir in Water. Antimicrobial Agents and Chemotherapy 59, 5196, doi:10.1128/AAC.00886-15 (2015).

30. Gillman, A. et al. Resistance mutation R292K is induced in influenza A(H6N2) virus by exposure of infected mallards to low levels of oseltamivir. PLoS One 8, e71230, doi:10.1371/journal.pone.0071230 (2013).

31. Gabriel, G. et al. The viral polymerase mediates adaptation of an avian influenza virus to a mammalian host. Proceedings of the National Academy of Sciences of the United States of America 102, 18590, doi:10.1073/pnas.0507415102 (2005).

32. Lazniewski, M., Dawson, W. K., Szczepińska, T. & Plewczynski, D. The structural variability of the influenza A hemagglutinin receptor-binding site. Briefings in Functional Genomics 17, 415–427, doi:10.1093/bfgp/elx042 (2017).

33. Bradley, K. C. et al. Analysis of Influenza Virus Hemagglutinin Receptor Binding Mutants with Limited Receptor Recognition Properties and Conditional Replication Characteristics. Journal of Virology 85, 12387, doi:10.1128/JVI.05570-11 (2011).

34. Lee, J. et al. Emergence of amantadine-resistant H3N2 avian influenza A virus in South Korea. Journal of clinical microbiology 46, 3788–3790, doi:10.1128/jcm.01427-08 (2008).

35. Evseev, D. & Magor, K. E. Innate Immune Responses to Avian Influenza Viruses in Ducks and Chickens. Vet Sci 6, 5, doi:10.3390/vetsci6010005 (2019).

36. Huang, Y.-F., Gulko, B. & Siepel, A. Fast, scalable prediction of deleterious noncoding variants from functional and population genomic data. Nat Genet 49, 618–624, doi:10.1038/ng.3810 (2017).

37. Younis, S. et al. Multiple nuclear-replicating viruses require the stress-induced protein ZC3H11A for efficient growth. Proceedings of the National Academy of Sciences 115, E3808, doi:10.1073/pnas.1722333115 (2018).

38. Byrd-Leotis, L. et al. Influenza binds phosphorylated glycans from human lung. Science Advances 5, eaav2554, doi:10.1126/sciadv.aav2554 (2019).

39. Byrd-Leotis, L. et al. Antigenic Pressure on H3N2 Influenza Virus Drift Strains Imposes Constraints on Binding to Sialylated Receptors but Not Phosphorylated Glycans. J Virol 93, doi:10.1128/jvi.01178-19 (2019).

40. van Dijk, J. G., Verhagen, J. H., Wille, M. & Waldenström, J. Host and virus ecology as determinants of influenza A virus transmission in wild birds. Current opinion in virology 28, 26–36, doi:10.1016/j.coviro.2017.10.006 (2018).

41. Latorre-Margalef, N. et al. Long-term variation in influenza A virus prevalence and subtype diversity in migratory mallards in northern Europe. *Proceedings*. Biological sciences 281, 20140098, doi:10.1098/rspb.2014.0098 (2014).

42. Jourdain, E. et al. Influenza virus in a natural host, the mallard: experimental infection data. PLoS One 5, e8935, doi:10.1371/journal.pone.0008935 (2010).

43. Pantin-Jackwood, M. J. et al. Pathogenicity and Transmission of H5 and H7 Highly Pathogenic Avian Influenza Viruses in Mallards. J Virol 90, 9967–9982, doi:10.1128/jvi.01165-16 (2016).

44. Keawcharoen, J. et al. Wild ducks as long-distance vectors of highly pathogenic avian influenza virus (H5N1). Emerging infectious diseases 14, 600–607, doi:10.3201/eid1404.071016 (2008).

45. Kleyheeg, E. et al. Deaths among Wild Birds during Highly Pathogenic Avian Influenza A(H5N8) Virus Outbreak, the Netherlands. Emerging infectious diseases 23, 2050–2054, doi:10.3201/eid2312.171086 (2017).

46. Wiethoelter, A. K., Beltran-Alcrudo, D., Kock, R. & Mor, S. M. Global trends in infectious diseases at the wildlife-livestock interface. Proc Natl Acad Sci U S A 112, 9662–9667, doi:10.1073/pnas.1422741112 (2015).

47. Webster, R. G., Bean, W. J., Gorman, O. T., Chambers, T. M. & Kawaoka, Y. Evolution and ecology of influenza A viruses. Microbiol Rev 56, 152–179 (1992).

48. Gill, F. D. D. (2019).

49. Wallensten, A. et al. Surveillance of influenza A virus in migratory waterfowl in northern Europe. Emerging infectious diseases 13, 404–411, doi:10.3201/eid1303.061130 (2007).

50. Wilcox, B. R. et al. Influenza-A viruses in ducks in northwestern Minnesota: fine scale spatial and temporal variation in prevalence and subtype diversity. PLoS One 6, e24010, doi:10.1371/journal.pone.0024010 (2011).

51. Ramey, A. M. & Reeves, A. B. Ecology of Influenza A Viruses in Wild Birds and Wetlands of Alaska. Avian Diseases 64, 109–122, 114 (2020).

52. Root, J. J., Shriner, S. A., Ellis, J. W., VanDalen, K. K. & Franklin, A. B. Transmission of H6N2 wild bird-origin influenza A virus among multiple bird species in a stacked-cage setting. Arch Virol 162, 2617–2624, doi:10.1007/s00705-017-3397-y (2017).

53. Verhagen, J. H. et al. Discordant detection of avian influenza virus subtypes in time and space between poultry and wild birds; Towards improvement of surveillance programs. PLoS One 12, e0173470, doi:10.1371/journal.pone.0173470 (2017).

54. Li, R. et al. Phylogeographic Dynamics of Influenza A(H9N2) Virus Crossing Egypt. Front Microbiol 11, 392, doi:10.3389/fmicb.2020.00392 (2020).

55. Helin, A. S. et al. A rapid and transient innate immune response to avian influenza infection in mallards. Molecular Immunology 95, 64–72, doi:https://doi.org/10.1016/j.molimm.2018.01.012 (2018).

56. Vanderven, H. A. et al. Avian influenza rapidly induces antiviral genes in duck lung and intestine. Mol Immunol 51, 316–324, doi:10.1016/j.molimm.2012.03.034 (2012).

57. Ito, T. et al. Molecular basis for the generation in pigs of influenza A viruses with pandemic potential. Journal of Virology 72, 7367–7373 (1998).

58. Bewley, C. A. Illuminating the switch in influenza viruses. Nature biotechnology 26, 60–62, doi:10.1038/nbt0108-60 (2008).

59. Gambaryan, A. S. et al. Receptor-binding profiles of H7 subtype influenza viruses in different host species. J Virol 86, 4370–4379, doi:10.1128/jvi.06959-11 (2012).

60. Tian, W. et al. The glycosylation design space for recombinant lysosomal replacement enzymes produced in CHO cells. Nature communications 10, 1785, doi:10.1038/s41467-019-09809-3 (2019).

61. Broszeit, F. et al. N-Glycolylneuraminic Acid as a Receptor for Influenza A Viruses. Cell reports 27, 3284–3294.e3286, doi:10.1016/j.celrep.2019.05.048 (2019).

62. Reed, L. J. & Muench, H. A SIMPLE METHOD OF ESTIMATING FIFTY PER CENT ENDPOINTS12. American Journal of Epidemiology 27, 493–497, doi:10.1093/oxfordjournals.aje.a118408 (1938).

63. Hoffmann, B. et al. New real-time reverse transcriptase polymerase chain reactions facilitate detection and differentiation of novel A/H1N1 influenza virus in porcine and human samples. Berliner und Munchener tierarztliche Wochenschrift 123, 286–292 (2010).

64. Pohlmann, A. et al. Swarm incursions of reassortants of highly pathogenic avian influenza virus strains H5N8 and H5N5, clade 2.3.4.4b, Germany, winter 2016/17. Sci Rep 8, 15, doi:10.1038/s41598-017-16936-8 (2018).

65. Moll, P., Ante, M., Seitz, A. & Reda, T. QuantSeq 3′ mRNA sequencing for RNA quantification. Nature Methods 11, i-iii, doi:10.1038/nmeth.f.376 (2014).

66. Andrews, S. FastQC: a quality control tool for high throughput sequence data. *Available:* https://www.bioinformatics.babraham.ac.uk/projects/fastqc/ *Accessed 2020 March 1* (2010).

67. Dobin, A. et al. STAR: ultrafast universal RNA-seq aligner. Bioinformatics 29, 15–21, doi:10.1093/bioinformatics/bts635 (2012).

68. Trapnell, C. et al. Transcript assembly and quantification by RNA-Seq reveals unannotated transcripts and isoform switching during cell differentiation. Nature biotechnology 28, 511–515, doi:10.1038/nbt.1621 (2010).

69. Love, M. I., Huber, W. & Anders, S. Moderated estimation of fold change and dispersion for RNA-seq data with DESeq2. Genome biology 15, 550, doi:10.1186/s13059-014-0550-8 (2014).

70. Blighe, K. EnhancedVolcano: Publication-ready volcano plots with enhanced colouring and labeling. Available: https://github.com/kevinblighe. (2018).

71. R: A language and environment for statistical computing v. 3.3.2 (R Foundation for Statistical Computing, Vienna, Austria, 2016).

72. Pertea, M. et al. StringTie enables improved reconstruction of a transcriptome from RNA-seq reads. Nature biotechnology 33, 290–295, doi:10.1038/nbt.3122 (2015).

73. Altschul, S. F., Gish, W., Miller, W., Myers, E. W. & Lipman, D. J. Basic local alignment search tool. Journal of molecular biology 215, 403–410, doi:10.1016/s0022-2836(05)80360-2 (1990).

74. Conway, J. R., Lex, A. & Gehlenborg, N. UpSetR: an R package for the visualization of intersecting sets and their properties. Bioinformatics (Oxford, England) 33, 2938–2940, doi:10.1093/bioinformatics/btx364 (2017).

75. R Core Team. R: A Language and Environment for Statistical Computing. R Foundation for Statistical Computing, Vienna, Austria. In: R Core Team, ed. Sincere Pumpkin Patch. 3.3.2 ed. (2016).

76. Wisniewski, J. R., Zougman, A., Nagaraj, N. & Mann, M. Universal sample preparation method for proteome analysis. Nat Methods 6, 359–362, doi:10.1038/nmeth.1322 (2009).

77. Parker, B. L. et al. Site-specific glycan-peptide analysis for determination of N-glycoproteome heterogeneity. Journal of proteome research 12, 5791–5800, doi:10.1021/pr400783j (2013).

78. Pett, C. et al. Effective Assignment of alpha2,3/alpha2,6-Sialic Acid Isomers by LC-MS/MS-Based Glycoproteomics. Angewandte Chemie (International ed. in English) 57, 9320–9324, doi:10.1002/anie.201803540 (2018).

79. Perez-Riverol, Y. et al. The PRIDE database and related tools and resources in 2019: improving support for quantification data. Nucleic acids research 47, D442–d450, doi:10.1093/nar/gky1106 (2019).

80. Varki, A. et al. Symbol Nomenclature for Graphical Representations of Glycans. Glycobiology 25, 1323–1324, doi:10.1093/glycob/cwv091 (2015).

